# Spatiotemporal Modeling of Mitochondrial Network Architecture

**DOI:** 10.1101/2024.01.24.577101

**Authors:** Keaton Holt, Julius Winter, Suliana Manley, Elena F. Koslover

## Abstract

In many cell types, mitochondria undergo extensive fusion and fission to form dynamic, responsive network structures that contribute to a number of homeostatic, metabolic, and signaling functions. The relationship between the dynamic interactions of individual mitochondrial units and the cell-scale network architecture remains an open area of study. In this work, we use coarse-grained simulations and approximate analytic models to establish how the network morphology is governed by local mechanical and kinetic parameters. The transition between fragmented structures and extensive networks is controlled by local fusion-to-fission ratios, network density, and geometric constraints. Similar fusion rate constants are found to account for the very different structures formed by mammalian networks (poised at the percolation transition) and well-connected budding yeast networks. Over a broad parameter range, the simulated network structures can be described by effective mean-field association constants that exhibit a non-linear dependence on the microscopic non-equilibrium fusion, fission, and transport rates. Intermediate fusion rates are shown to result in the highest rates of network remodeling, with mammalian mitochondrial networks situated in a regime of high turnover. Our modeling framework helps to elucidate how local parameters that govern mitochondrial interactions give rise to spatially resolved dynamic network structures at the cellular scale.

## I. INTRODUCTION

Eukaryotic cells rely on mitochondria, pleomorphic organelles specialized for ATP production, to supply their metabolic needs. In addition to their role as an energetic powerhouse, mitochondria participate in a broad array of cellular functions, including Ca^2+^ homeostasis, lipid storage and production, and programmed cell death [1]. In recent decades, the physical structure of mitochondria has been found to be vitally intertwined with cellular function, state, and health [2–5]. Across a variety of cell types, mitochondria are observed to form dynamic, interconnected spatial networks which rearrange on the order of seconds [6] through fusion and fission of mitochondrial membranes, and motion of the organelles through the intracellular space [7–9]. These networks can exhibit a variety of structures, including three-dimensional partially fragmented architectures in mammalian cells [6], and cortical surface-bound extensively-connected networks in yeast cells [10].

The formation of interconnected mitochondrial networks is implicated in a host of cellular functions [11]. These include protection against stress and perturbations to membrane potential [12], modulation of ATP production rate and efficiency [13], and regulation of calcium signaling [14]. The networked mitochondrial architecture is also thought to play an important role in quality control [3, 11, 15]. Mitochondrial proteins and DNA are particularly prone to oxidative damage as a result of high respiration rates, with the mutation rate of human mtDNA estimated as roughly 400-fold higher than in the nucleus [16]. Fusion and fission enable mitochondria to share their contents [17], allowing for potential complementation of damaged components [18, 19]. In addition, dynamic remodeling of the mitochondrial network, together with selective fusion [2], fission [20], and/or autophagy [8, 21], makes it possible to segregate and recycle damaged mitochondria, improving the health of the overall population.

When quality control is dysfunctional, accumulated damage to mtDNA and mitochondrial proteins over multiple cell cycles leads to loss of mitochondrial function, a decline which is associated with ageing [22]. Decreased mitochondrial function, coupled with modification of network structure, is also associated with neurological diseases such as Alzheimer’s [23], Parkinson’s [24], and Huntington’s Disease [25]. Fragmented mitochondrial networks are frequently observed in each of these pathologies, suggesting impairment of the fusion and fission machinery [26]. Indeed, induced overexpression of fusion proteins or repression of fission proteins may rescue the normal phenotype from an initially diseased cell state [27].

Mitochondrial network structure can be altered by genetic and pharmacological interventions as well as nutrient and environmental conditions. The upregulation or overexpression of mitochondrial fusion proteins (MFN1, MFN2, OPA1 in mammalian cells, and Fzo1, Mgm1 in yeast [3]) can lead to hyperfused, highly-connected networks [28], and may counteract pathological fragmentation [27]. Genetic disruption of fission proteins [29] (DRP1, Dnm1 in mammalian and yeast cells, respectively [3]) also leads to mitochondrial elongation, while treatment with toxins such as oligomycin [6], FCCP [30], and paraquat [28] induces fragmentation of the network. Nutrient availability has been shown to regulate mitochondrial network volume [31] and structure. Cells grown in excess glucose undergo mitochondrial fragmentation [13], while those in a state of nutrient deprivation exhibit increased mitochondrial tubulation and connectivity [32]. Such tubulation is generally accompanied by altered intra-mitochondrial structure, with a higher density of cristae junctions on the inner membrane leading to increased ATP synthesis [4, 33].

Recent advances in imaging technology have allowed for real-time tracking and quantification of mitochondrial network structure [6, 10, 34]. However, our understanding of the physical processes and principles that govern these dynamic structures remains incomplete. Previous models have largely relied on aspatial [35], continuum [36], lattice-based [11], or two-dimensional [8, 37] approaches. This prior work elucidated the role of effective cellular-scale fusion and fission rates [28, 35], and the ability of networks to distribute or segregate mitochondrial contents [8, 11, 37]. Nevertheless, the impact of mitochondrial mechanics, local interaction kinetics, and transport dynamics on network structure remains opaque.

The fusion and fission of mitochondrial units is in many ways analogous to reversible clustering in molecular-scale systems. Past work on molecular aggregation has examined the structures formed by interacting monomers undergoing Brownian motion. In models of reversible aggregation, the probabilities to form and break bonds between monomers control the properties of the resulting structure [38]. Other formulations rely on a temperature-dependent short-range attraction between colliding monomers to hold the structure together [39]. Above a critical value of the bonding probability or attraction strength, called the percolation threshold, most particles will be members of a single supercluster [40, 41]. The percolation threshold has been extensively studied for systems of various constituent particle shapes such as spheres [42], ellipsoids [43], and spherocylinders [44]. Other studies have examined the role of particle mobility on structure [45, 46], finding that increased mobility leads to a higher degree of compactness. Notably, mitochondrial networks differ from many previously studied clustering systems in that their fusion and fission processes are energy-driven and thus out of thermal equilibrium. This energy consumption has the potential to break some of the thermodynamic constraints linking mitochondrial motility, flexibility, and the configuration-dependent fusion/fission balance, thereby altering the relationship between network morphology and microscopic dynamics.

Here we ask how local interactions between individual pairs of mitochondria, together with the motion of the organelles through space, gives rise to emergent large-scale network structure. In particular, we elucidate the relationship between microscopic parameters such as the fusion rate for adjacent mitochondria, the bending stiffness of the mitochondrial tubules, and the mobility of individual mitochondria, with macroscopic parameters such as the segment length and connectivity of the resulting network. We find that network structures below or near percolation can be approximately predicted from microscopic rate constants, in both 3D and surface-bound systems and demonstrate that intermediate fusion rates allow for the most rapid rearrangement of network structure. These results lay the groundwork for developing a fundamental mechanistic understanding of how diverse experimental perturbations to mitochondrial morphogens modulate the overall dynamics, and thereby the health and function, of the network.

## II. MODEL DESCRIPTION

We model the mitochondrial network using a 3D, dynamic, graph-based formulation in which edges represent discrete mitochondrial units of ground-state length *l*_0_ = 0.5*μ*m and steric radius *r*_s_ = 0.1*μ*m. Nodes represent the endpoints of each mitochondrion and serve as the locations where forces are applied and fusion/fission events can occur. The positions of the nodes evolve over time according to an overdamped Langevin equation:

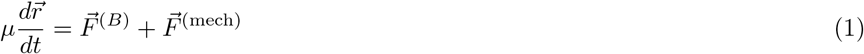

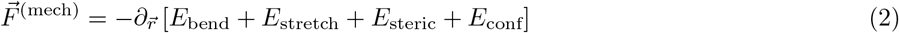

with *μ* the friction coefficient for each node, 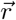 the vector of node positions and 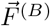 and 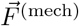 corresponding to the Brownian and mechanical forces, respectively. The stochastic Brownian forces are uncorrelated in time: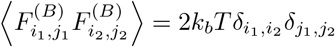, where *i* refers to the node index, *j* to the spatial dimension, and *k*_*b*_*T* is the thermal energy.

The bending energy is defined by:

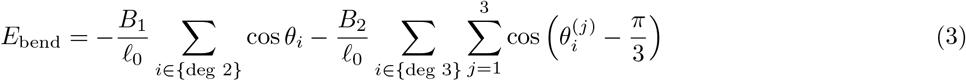

where the first term depends on the deviation of each degree-2 node from a straight configuration and the second term depends on the deviation of each angle in a degree-3 junction from its ground-state value of 60^*°*^. The mechanical energy also includes quadratic penalties for each edge length deviating from its ground state value (*E*_stretch_), for two non-connected edges approaching closer than the minimal steric distance (*E*_steric_), and for any node escaping a spherical confinement (*E*_conf_). Details of the energetic terms, the choice of default parameters, and the time integration of Eq. 1 are given in Methods.

Topological remodeling of the network is implemented via local fusion and fission events at the nodes (Fig. 1A,B). We assume that fusion may only occur between a pair of degree-1 nodes (tip-tip fusion) or between a degree-1 and a degree-2 node (tip-side fusion). This limits the network to nodes of degree 1-3, in keeping with prior work showing that higher-degree nodes are rarely observed in mitochondrial networks [28, 35]. Fusion is allowed when the nodes are separated by a distance *≤* 2*r*_*c*_, where *r*_*c*_ is the contact radius. The fusion rates are taken to be dependent on the orientation of the fusing edges, precluding fusion events that lead to very high bending energies. The implicit assumption is that fusion machinery on the mitochondrial surface can only perform its function if the mitochondrial segments are favorably aligned. By default, we make the fusion rates for tip-tip and tip-side fusion 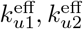, respectively) proportional to a Boltzmann factor incorporating the relevant bending energies:

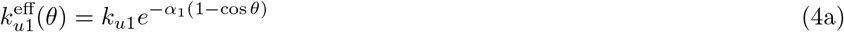

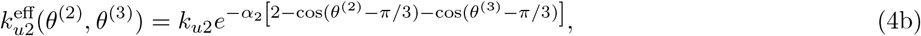

where *θ* is the angle formed by the edges meeting at the new degree-2 node and *θ*^(2)^, *θ*^(3)^ are the angles formed by the incoming 3rd edge relative to the existing edges making up the former degree-2 junction. As with the mechanical bending at the 3-way junction, angles of *π/*3 are assumed to be preferred for fusion. However, for degree-3 junction formation, only the two angles involving the newly added segment are used to define the fusion rate. The parameters *k*_*ui*_ set the fusion rate when all edges are perfectly aligned, while *α*_*i*_ sets the sensitivity of the fusion machinery to deviations from the preferred junction angles.

**FIG. 1.**
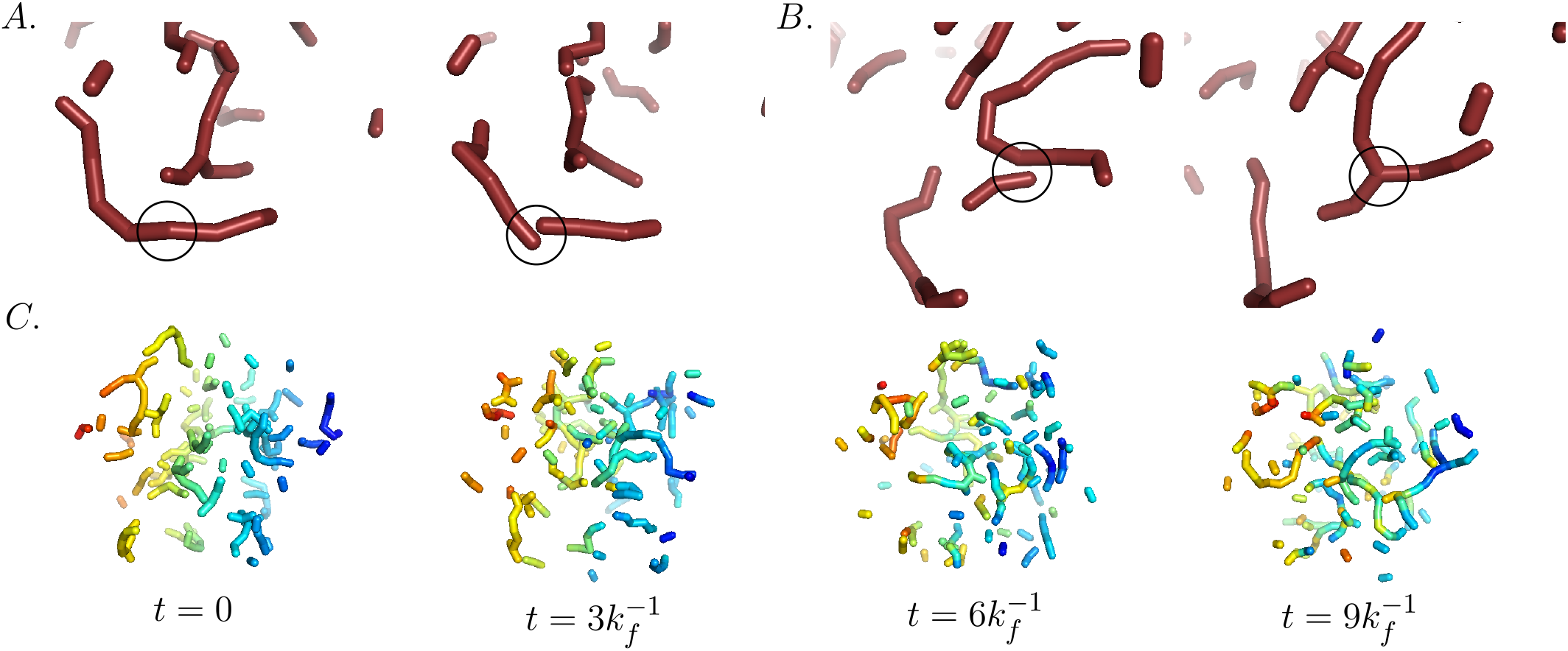
Fission and fusion reshape the mitochondrial network. **A**. An example of tip-tip fission at a degree-2 node. **B**. An example of tip-side fusion to form a degree-3 junction. **C**. Snapshots of an example mitochondrial network simulation. Edges are colored according to their initial position to visualize mixing.

The angular dependence of fusion rates has not been empirically established. The form selected here satisfies the general principle of permitting fusion only when it will not lead to large energy penalties at the newly-formed junctions. The default angle-sensitivities are set to *α*_*i*_ = *B*_*i*_*/*(*R*_0_*k*_*b*_*T*). For the degree-2 case this is the expected behavior for a system in thermal equilibrium, where fusion and fission obey detailed balance. For the degree-3 case, detailed balance is broken because the fusion rate excludes the change in energy associated with the pre-existing junction angle.

Fission occurs at non-terminal network nodes, with a rate of *k*_f_ for degree-2 nodes and 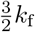 for degree-3 junctions. In keeping with prior models [35], these rates assume that the fission machinery docks on the edges surrounding the junction, so that fission is proportionally more likely to occur at a junction with 3 adjacent edges. Following fission of a degree-2 node, the tips of the resulting segments (defined as one steric radius beyond the new degree-1 node positions) meet at the former node location. Upon fission at a degree-3 junction, one edge is selected at random to break from the others. The newly formed tip of this edge is placed at the former junction while the remaining degree-2 node is moved away in the opposite direction from the original junction by a distance *r*_s_ (the steric radius). This approach ensures that a very rapid fission and re-fusion event would result in no net change in the mechanical energetics at the junction. Details of the fusion and fission processes are illustrated in Supplemental Material.

The dynamic model is evolved forward in time using Brownian Dynamics simulations, with the appropriate probabilities of fusion and fission at each time-step based on the remodeling rates defined above. The simulations are carried out in dimensionless units, with the length scale set by 2*R*_0_ = 1, the time scale by 1*/k*_*f*_ = 1 (where *k*_*f*_ is a fission rate, discussed below), and the force scale by *μ* = 1.

To convert to real units, we estimate *k*_*f*_ *≈* 0.5min^*−*1^. As discussed in the Results, this gives an overall rate of ‘complete fission’ (not followed by re-fusion to the same node) on the order of 0.1 per minute per mitochondrion, consistent with prior experimental and modeling studies [2, 8, 47–49]. The default effective temperature is set to *k*_*b*_*T* = 1 in dimensionless units, yielding a diffusivity for an isolated mitochondrial edge-unit of *D*_1_ = 0.25*μ*m^2^*/*min, consistent with measurements of stochastic mitochondrial motion within mammalian cells [6, 50, 51].

The resulting network configurations rearrange over time-scales governed by the fusion and fission rates as well as the effectively diffusive movement of the mitochondrial units. Substantial rearrangements of network topology occur over a time period of a few minutes (Fig. 1C).

## III. RESULTS

### A. Network architecture emerges from the balance of fusion and fission rates

As with molecular clustering systems that undergo reversible aggregation [38, 52], the steady-state structure of the mitochondrial network depends on the ratio of local fusion and fission rate parameters *k*_*ui*_*/k*_*f*_ (Fig. 2). For a fixed fission rate, and low fusion, the mitochondria are largely fragmented into small disjoint segments. At higher fusion, networks become well-connected, containing numerous degree-2 nodes and degree-3 junctions. When the fusion rates are similar (*k*_*u*2_ = *k*_*u*1_), degree-3 junctions dominate within well-connected networks (Fig. 2A, left). By contrast, when the tip-side fusion rate is reduced (*k*_*u*2_ = 0.1*k*_*u*1_), there is a broad intermediate regime with long linear chains containing many degree-2 nodes and only a small number of degree-3 junctions (Fig. 2A, right).

**FIG. 2.**
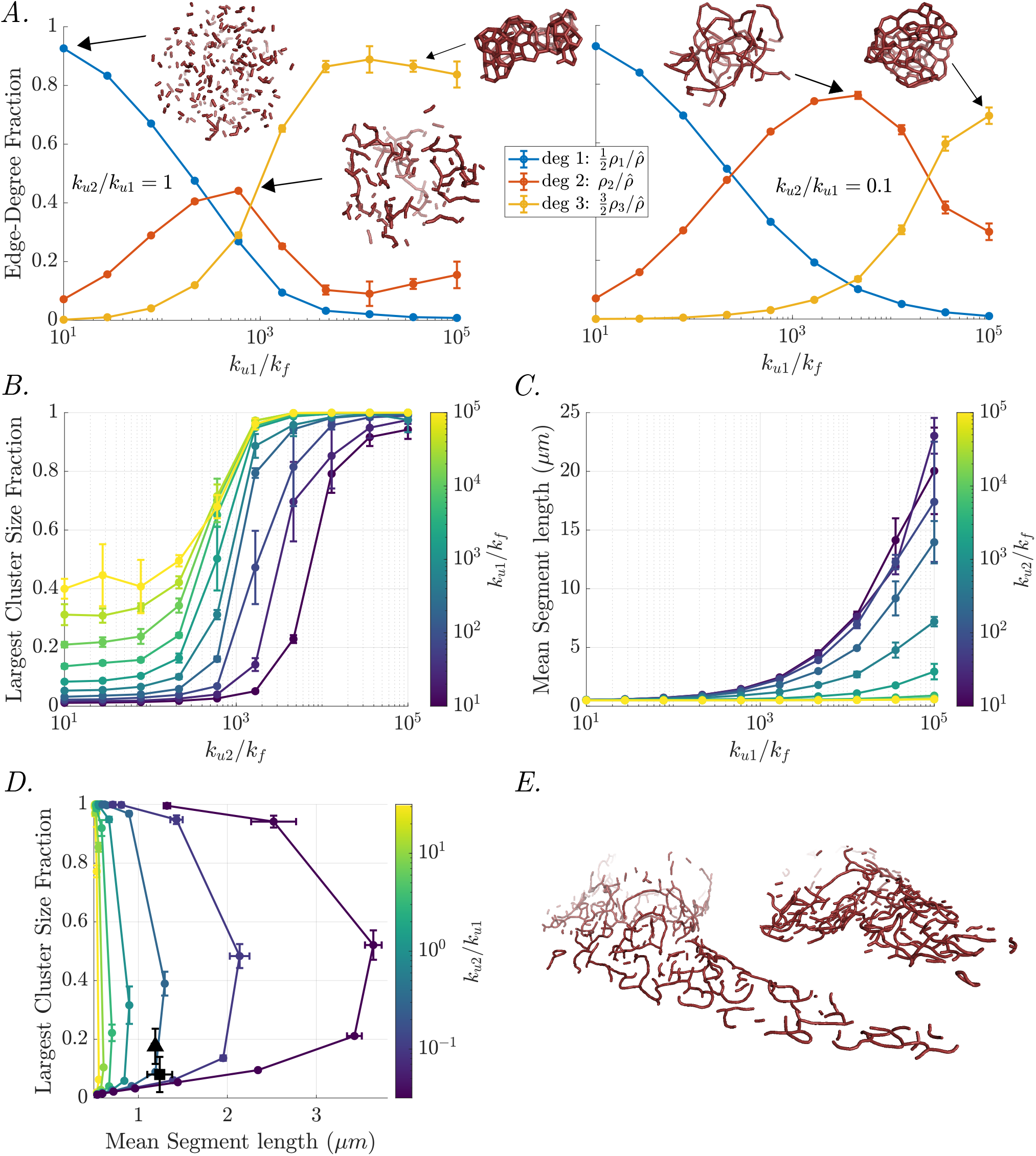
Fusion-fission rate balance principally determines network structure and connectivity. **A**. The fraction of edges connected to nodes of each degree (1-3) in the network is plotted as a function of increasing fusion rate. Insets show example networks. Left: *k*_*u*2_ = *k*_*u*1_, Right: *k*_*u*2_ = 0.1*k*_*u*1_. **B**. The largest cluster size fraction ⟨*L*_*G*_ ⟩*/L* is plotted as a function of increasing *k*_*u*2_ for fixed values of *k*_*u*1_ (colors). **C**. The mean segment length ⟨*L*_seg_ ⟩is plotted as a function of increasing *k*_*u*1_ for fixed values of *k*_*u*2_ (colors). **D**. Simulated (colored curves) and experimental (black triangle: hiPSC cell [6], black square: COS-7 cells) networks are plotted in the space of largest cluster size fraction and mean segment length. Colors indicate fixed values of the ratio *k*_*u*2_*/k*_*u*1_ while dots represent specific values of *k*_*u*1_, which increases from low to high cluster size along each curve. Error bars indicate the standard deviation across simulation trials (A-D) or experimental images (D). **E**. Examples of experimentally derived network structures from a COS-7 cell (left) and a hiPSC cell [6] (right).

In keeping with prior measurements of mitochondrial networks [10, 28, 35], we quantify the network architecture using two experimentally accessible metrics. The largest cluster size fraction ⟨*L*_*G*_*/L*⟩ is defined as the mitochondrial length in the largest connected component of the network, normalized by the total mitochondrial length. This metric quantifies the overall network connectivity, with a value of 1 indicating a fully-connected structure and low values corresponding to fragmented populations. As the tip-side fusion rate *k*_*u*2_ is dialed up, the system undergoes a percolation transition with network fragments coalescing into a single super-cluster. This is indicated by the sharp jump in ⟨*L*_*G*_*/L*⟩ (Fig. 2B). Notably, the onset of percolation is shifted to higher values of *k*_*u*2_ when the rate of tip-tip fusion is reduced, due to the ensuing scarcity of degree-2 nodes necessary for forming junctions. Near the percolation transition, small changes in the fusion rate can greatly alter the network architecture. Previous studies have indicated that mammalian mitochondrial networks tend to balance near this percolation threshold [28, 35, 53], allowing for profound alterations in network morphology in response to small environmental changes.

A separate structural metric is the mean segment length ⟨*L*_seg_ ⟩= *𝓁*_0_*N*_edge_*/N*_seg_, where a segment is defined as a linear chain of mitochondrial units, terminating at either a degree-1 node or a degree-3 junction. The number of segments can be computed as *N*_seg_ = *N*_edge_ *N*_degree-2 nodes_ + *N*_free loops_, where a free loop is a connected component containing only degree-2 nodes. A large value of ⟨*L*_seg_ ⟩indicates the presence of numerous extended linear mitochondria with few branches. Notably, a maximally-connected network (containing entirely branching nodes) and a completely fragmented network (only degree-1 nodes) have the same mean segment length (unit length *𝓁*_0_ = 0.5*μ*m). High values of ⟨*L*_seg_⟩ require tip-tip fusions to dominate, with *k*_*u*1_ *≫ k*_*u*2_ (Fig. 2C).

By varying both microscopic fusion rates (relative to a fixed fission rate *k*_*f*_), a broad region of the two structural features (⟨*L*_*G*_ ⟩*/L* and ⟨*L*_seg_ ⟩) can be accessed (Fig. 2D). For a given largest cluster size, the mean segment length can be tuned by altering the ratio of tip-side versus tip-tip fusion (*k*_*u*2_*/k*_*u*1_). High values of this ratio (yellow curve) give networks with very short segment lengths across the entire spectrum of connectivity. Low values of the ratio (purple curve) allow for long segment lengths throughout an intermediate regime where networks are neither fully fragmented nor fully percolated.

We identify the experimentally relevant fusion parameters by comparing with published mitochondrial network structures from a set of 93 images of a human induced pluripotent stem cell (hiPSC, provided in Ref. [6]) and from 37 images of COS-7 cells grown on patterns (obtained as described in Methods). These networks (Fig. 2E) have mean segment lengths of ⟨*L*_seg_⟩ *≈* 1.19, 1.24*μ*m and largest cluster fraction ⟨*L*_*G*_*/L*⟩ *≈* 0.18, 0.08, respectively. Both mammalian cell types yield consistent estimates of the relative fusion ratio: *k*_*u*2_*/k*_*u*1_ *≈* 0.37 *±* 0.06, and 0.26 *±* 0.13 respectively.

The tip-tip versus tip-side microscopic fusion rate constants for spatially proximal and orientationally well-aligned segments are thus expected to be quite similar in magnitude. This estimate is in stark contrast to the overall global association constants predicted in prior work, which indicated a global tip-side fusion rate that was lower by a factor of 100 [28] to 10000 [35] when compared to tip-tip fusion.

### B. Spatial constraints and local non-equilibrium kinetics modulate network structure

In order to map from microscopic parameters to large-scale network architecture, we first consider a simplified mean-field model of bulk kinetics. Mass action models have been employed in a wide variety of contexts to determine the behavior of molecular-scale systems from a set of binding and unbinding rates [54–56]. These aspatial models rely on two key assumptions: a well-mixed system (spatially uniform reactant concentrations) and reaction rates that are directly proportional to concentrations. The steady-state concentrations are then found by defining the rate of change for each component and setting the resulting equations to zero.

A mass action model for mitochondrial networks, originally developed by Sukhorukov *et. al*. [35], treats nodes of different degree (*X*_1_, *X*_2_, *X*_3_) as distinct species, undergoing the following reactions:

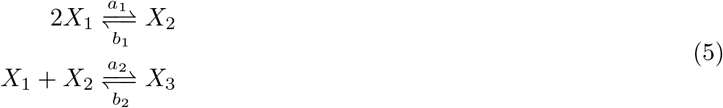

where *a*_1_, *a*_2_ represent bulk rate constants for tip-tip fusion and tip-side fusion, and *b*_1_, *b*_2_ are the fission rates for degree-2 and degree-3 nodes. These bulk rate constants are effective parameters that depend not only on the microscopic fusion rate between a pair of nearby nodes but also on the frequency with which nodes come into contact with each other in an appropriate orientation for fusion to occur.

The steady-state equations expressed in terms of the concentration of nodes (*ρ*_1_, *ρ*_2_, *ρ*_3_ for degree 1, 2, and 3, respectively) are then given by:

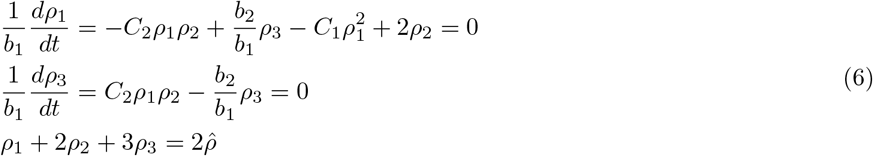

where the third equation ensures mass conservation and 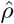 is the total concentration of edge units in the network. *C*_1_ = *a*_1_*/b*_1_ and *C*_2_ = *a*_2_*/b*_1_ are the effective association constants for the fusion/fission reactions, with dimensions of inverse concentration. These association constants are directly related to the steady-state densities of nodes according to: 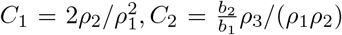. They are intensive properties of the system, so their values remain constant if the size of the entire system is altered while maintaining the same node densities. The quantities 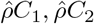 are dimensionless values that describe the overall steady-state balance between fragmentation and connectivity for tip-tip and tip-side fusions, respectively.

As shown in Fig. 3, the association constants for degree-2 and 3 fusions vary as a function of the corresponding microscopic fusion rate. The edge-unit densities and node counts of different degrees can be directly estimated for experimental data (details in Methods), allowing the extraction of individual microscopic fusion to fission ratios. For the two sets of mammalian mitochondrial networks considered here, we estimate *k*_*u*1_*/k*_*f*_ = (1100 *±* 100), (1200 *±* 600) and *k*_*u*2_*/k*_*f*_ = (470 *±* 40), (400 *±* 200), giving a similar ratio (*k*_*u*2_*/k*_*u*1_) to that extracted from measurements of segment length and largest cluster size (Fig. 2).

**FIG. 3.**
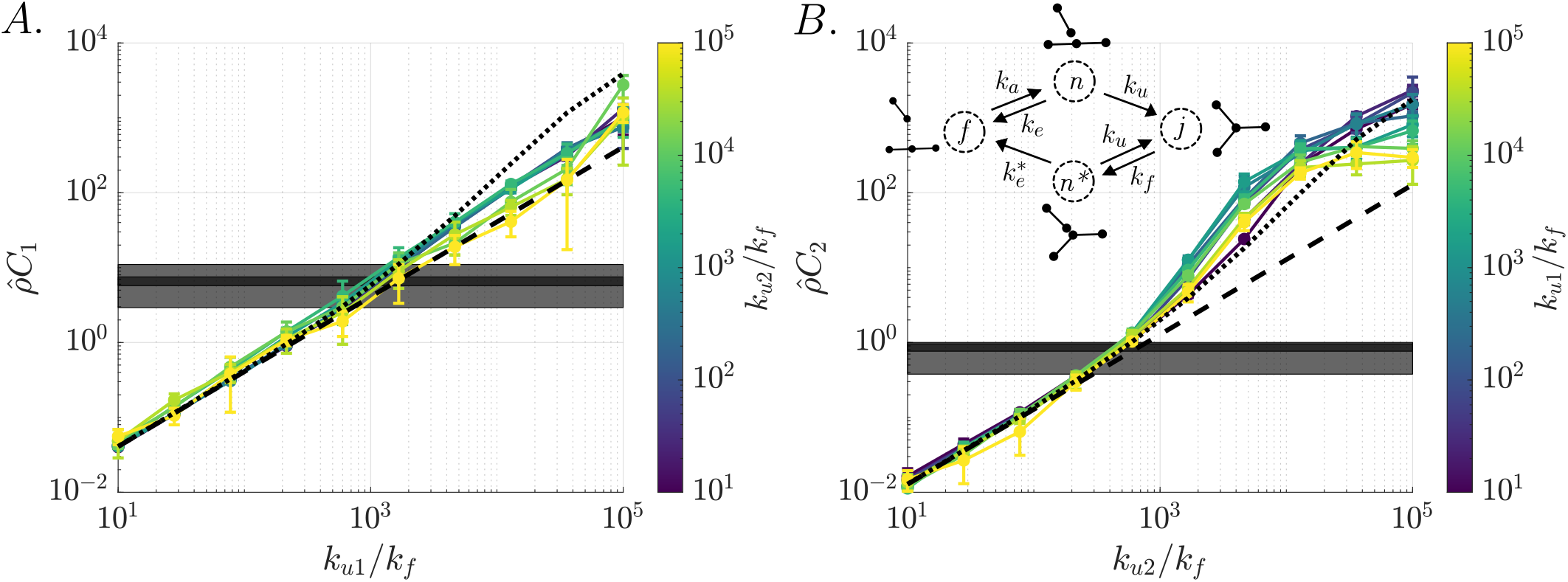
Mapping local parameters to effective association constants. **A**. The tip-tip association constant, *C*_1_, is plotted as a function of the local fusion parameter *k*_*u*1_ for different values of *k*_*u*2_ (colored lines). **B**. The tip-side association constant, *C*_2_, is plotted as a function of the local fusion parameter *k*_*u*2_ for different values of *k*_*u*1_ (colored lines). Inset: a simplified kinetic model approximates the deviation in *C*_2_ from equilibrium at intermediate fusion rates. In A, B dashed lines show the equilibrium mean-field prediction 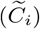. Dotted lines show predictions from the simplified kinetic model (*Ĉ*_*i*_). Error bars show standard deviation across simulation trials. Gray boxes show the mean and standard deviation for experimental networks from hiPSC [6] (dark gray) and COS-7 cells (light gray).

In our spatially-resolved model, fusion between mitochondria can only occur when two nodes are sufficiently close to each other and the connecting edges are favorably oriented. We estimate the effective bulk fusion rate according to 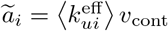, where *v*_cont_ is the contact volume surrounding a node and 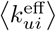 is the local fusion rate averaged over possible edge orientations within the contact volume (see Methods for details). The predicted association constants are then given by 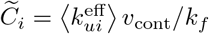

The mean-field estimates of the association constants are accurate for low fusion-to-fission ratios, where the networks are largely fragmented. However, at intermediate fusion (near the percolation transition), the observed tip-side association constants scale super-linearly with the fusion rate constant, and at high fusion (fully percolated networks), they flatten out to a sub-linear scaling. Both discrepancies indicate that the mitochondrial network node associations cannot be taken to obey the well-mixed assumptions of bulk kinetics.

Whenever a fission event occurs, the newly distinct nodes are located close to each other and are capable of rapidly re-fusing before they have a chance to explore the entire domain. For degree-3 nodes in particular, the distribution of edge orientations after fission differs from the distribution when a segment approaches for the bulk. Fission leaves the remaining degree-2 node in a strained, more highly bent state compared to its equilibrium distribution. Further, newly separated nodes are always placed at minimal steric separation, 2*r*_*s*_, whereas fusion can occur over a range of separations out to the maximal contact separation, 2*r*_*c*_ (see Supplemental Material for details). Overall, the state of a pair of edges immediately after fission cannot be treated as identical to the state immediately before fusion, breaking the assumptions of the mean-field model.

We can build a simplified kinetic model encompassing this effect by defining a set of discrete states for a pair of nodes (Fig. 3B, inset). The *f* state describes two nodes far apart in the domain, the *n* state describes the pair within contact range after finding each other from a distance, the *n*^*∗*^ state has the pair within contact range just after a fission, and the *j* state has the pair joined together into a single node. Notably, the *n*^*∗*^ state tends to favorably align recently-separated nodes for re-fusion, whereas bulk nodes tend to be less-favorably aligned. This system of states contains one-way arrows and thus fails to obey detailed balance [57, 58]. So long as the two contact states *n* and *n*^*∗*^ have different rates of escape to the far *f* state, the system is not in thermodynamic equilibrium. It should be noted that the biomolecular machinery responsible for mitochondrial fusion and fission consumes GTP [59], and the fission-fusion cycle is thus expected to be a non-equilibrium process. In this sense, the mitochondrial dynamics fundamentally differ from classic molecular clustering systems which have previously been explored with spatial simulations [38, 39, 52].

Solving for the steady-state probabilities of the abstract model system in Fig. 3B inset (see Methods for details) gives the effective association constant 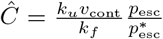, where 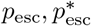 are the probabilities of escape from the *n, n*^*∗*^ state, respectively, into the *f* state before fusion. For the mitochondrial dynamics, fission results in two edges within steric contact and in more favorable orientations than would be expected at equilibrium, so that 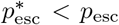. The escape probabilities for the two states can be estimated by computing the diffusive escape out of contact range from a spherical shell domain in the presence of a spatially homogeneous reaction rate, as described in Supplemental Material.

The effective association constants,

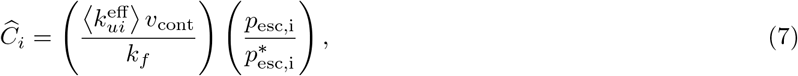

approximately match those extracted from simulations for low and intermediate junction fusion rates (Fig. 3A,B).

At low fusion rates, the escape probabilities approach 1 and the association constants depend linearly on the corresponding microscopic fusion rate. At intermediate fusion rates, the reduced probability of escaping implies likely re-fusion following each fission event, leading to enhanced association when compared to an equilibrium system. The effect is most clearly seen for the tip-side association constant *C*_2_ measured near the percolation transition. Here, the simplified model approximately matches simulation results in the limit of short segment lengths (low *k*_*u*1_, purple curve), where the network is partially fragmented into disjoint triskelion structures. Higher values of tip-tip fusion (green curve) increase the association constant still further, possibly by bringing the danging nodes of nearby triskelions in closer spatial proximity to each other. By contrast, very high values of *k*_*u*1_ (yellow curve) decrease the junction association constant in the intermediate regime, by forming very long linear segments between junctions that greatly slow the ability of dangling ends to find each other.

At very high tip-side fusion rates, the system collapses into a fully percolated network. Even in this regime, however, there are still some degree-1 and 2 nodes remaining at the network periphery. As a direct consequence of spatial and mechanical constraints, these nodes cannot reach each other to undergo fusion. This results in a saturation of the association constant *C*_2_ at high microscopic *k*_*u*2_.

For the tip-tip association constant *C*_1_, there is a smaller deviation from the linear equilibrium behavior. In contrast to the out-of-equilibrium orientation of segments upon tip-side fission, a tip-tip fission event places the newly formed segments into an orientational distribution proportional to the fusion rate, as expected for detailed balance. The only non-equilibrium effect arises from the close positioning of nodes upon fission. At very high rates of the microscopic tip-tip fusion parameter *k*_*u*1_, mitochondria associate into highly connected, linear structures whose distant peripheral ends are incapable of finding each other within the timescale of the simulation. The association constant thus drops below the values predicted by the approximate model (Eq. 7).

Our simulations highlight the importance of spatial locality and constraints in determining the steady-state structure of the mitochondrial network. Beyond the fragmented regime, details of the non-equilibrium fission and fusion process can substantially alter the network connectivity away from the mass-action prediction. The mass-action model provides a useful framework for computing structural metrics of network architecture based on two effective large-scale association parameters (*C*_1_, *C*_2_) [28, 35]. The more detailed spatial model described here goes one step further in estimating these effective parameters from the microscopic fusion and fission rates as well as the mechanical properties of the mitochondria.

### C. Surface-bound network architectures exhibit higher connectivity

Our model for mitochondrial network structure can be applied to other cell types and geometries, such as the budding yeast *S*.*cerevisiae*. The mitochondria of these small (*∼* 5*μ*m in diameter) cells are bound to the inner surface of the cell membrane, so that mitochondrial units are effectively constrained to move on a 2D shell. Notably, yeast mitochondrial networks are known to be more highly connected than mammalian networks, well beyond the percolation transition [10]. By incorporating a tethering force that holds mitochondria near a spherical boundary, we can generate surface-bound network structures (Fig. 4A) and estimate the underlying microscopic parameters that account for the observed network connectivity.

**FIG. 4.**
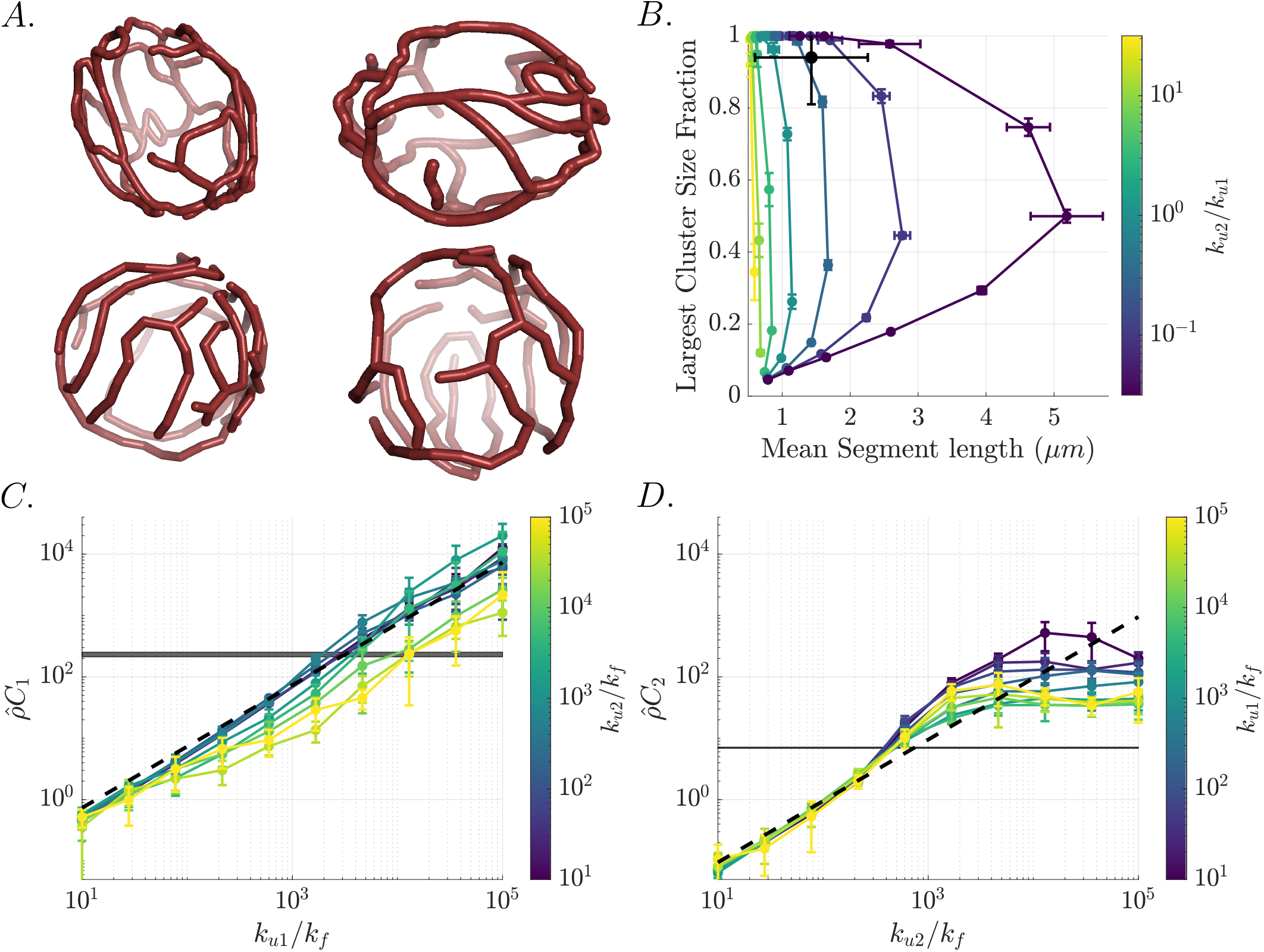
Structure of surface-bound mitochondrial networks. **A**. Example yeast networks obtained from live-cell imaging (top, from [10]) and via simulation (bottom). **B**. The largest cluster size fraction ⟨*L*_*G*_ *⟩/L* and mean segment length ⟨*L*_seg_ ⟩are plotted for different values of the ratio *k*_*u*2_*/k*_*u*1_ (colored curves). The black point represents the mean and standard deviation of 2794 experimental networks from [10]. **C**. The tip-tip association constant, *C*_1_, is plotted as a function of the local fusion parameter *k*_*u*1_ for different values of *k*_*u*2_ (colored lines). **D**. The tip-side association constant, *C*_2_, is plotted as a function of the local fusion parameter *k*_*u*2_ for different values of *k*_*u*1_ (colored lines). The black dashed curves show our analytic predictions from the equilibrium mean-field model; gray boxes highlight the mean and standard error of the extracted values from [10]. Simulations are performed in a spherical cell of radius *R* = 2.3*μ*m, with total mitochondrial length 60*μ*m to approximate experimentally observed yeast networks. Error bars in B-D show standard deviation over simulation trials.

As in the case of 3D networks, varying the ratio of tip-side versus tip-tip fusion rates, as well as the fusion to fission ratios, enables the generation of a family of structures with different mean segment lengths and largest cluster size fractions (Fig. 4B). We extract these two metrics from a previously published database of visualized yeast network structures [10], and compare to the simulation parameter sweeps. Experimental networks lie in the neighborhood of *k*_*u*2_*/k*_*u*1_ = 0.30 *±* 0.02 (mean *±* standard error), similar to the ratios obtained for 3D networks. This correspondence between microscopic parameter ratios across networks in different cell types suggests that the balance between tip-side and tip-tip fusions is a robust fundamental property of mitochondrial geometry and/or fusion machinery.

As in the case of 3D mammalian cell networks, we can turn to the simplified aspatial model (Eq. 6) to map our microscopic parameters to large-scale network structure. In a 2D system, the effective bulk fusion rate becomes a function of the contact area *a*_cont_ rather than the contact volume, and the local fusion rate is averaged over the possible 2D edge-unit orientations in this contact area. The node concentrations take on units of 1/area. With these modifications we can write the effective association constants: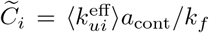. As shown in Fig. 4C-D, the mean-field approximation matches simulation results in the low-fusion regime. Intermediate and high values of tip-side fusion lead first to enhanced values of *C*_2_ (due to rapid refusion) and then to a flattening below the mean-field prediction (due to mechanical constraints on node accessibility), analogous to the 3D networks. Notably, the saturation at high fusion occurs for values of 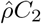 that are about an order of magnitude lower than in the 3D network structures. This implies that tethering to a surface makes it more difficult for remaining unfused nodes to reach each other, leading to frustrated network structures with limited maximal connectivity.

For any given set of microscopic fusion-to-fission ratios *k*_*ui*_*/k*_*f*_, the simulated surface-bound networks are more highly connected (higher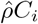) than their 3D counterparts in Fig. 3. This effect arises from the effectively higher density of nodes when networks are confined to a thin shell next to the cell boundary. Extracting individual parameter values from the empirical yeast network database (Fig. 4C, D, gray line) yields *k*_*u*1_*/k*_*f*_ = 1900 *±* 100, *k*_*u*2_*/k*_*f*_ = 480 *±* 10 (mean *±* standard error). These rates are only slightly higher than the mammalian cell estimates. If these microscopic fusion rates are applied to 3D networks with the appropriate unit-edge density for mammalian cells, they yield values of 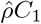 and 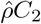 that are within the range observed for mammalian mitochondrial networks (see Fig. 3A,B). For surface-bound networks, the values of 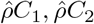 are approximately an order of magnitude higher (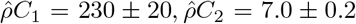 in yeast cells versus 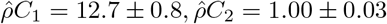 in mammalian cells). We therefore conclude that the higher connectivity observed in yeast networks is a result of geometric constraints that confine mitochondria to the cell surface, and the concomitant rise in mitochondrial density at the boundary, rather than substantial alterations in the balance of microscopic fusion-to-fission rates.

### D. Mitochondrial flexibility and orientational fusion sensitivity modulate network connectivity

In this model, the angular alignment of mitochondrial units plays two separate roles in determining network structure. First, the bending moduli *B*_1_, *B*_2_ control the stiffness of fused degree-2 and degree-3 junctions, respectively. Second, the parameters *α*_1_, *α*_2_ set the angular sensitivity of the fusion rate for forming each type of junction. The overall local fusion rate scales in proportion to the Boltzmann factor: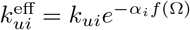, where *f* (Ω) is a function of the segment orientations, as in Eq. 4. By default, we set *α*_*i*_ = *B*_*i*_*/𝓁*_0_*k*_*b*_*T*, restricting fusion events to occur only for angular configurations that will give a low mechanical energy upon fusion. This choice of angle-dependence relates the fusion rate for a given edge orientation to the equilibrium probability of that orientation just before a fission event, with direct proportionality between the two for degree-2 junctions. Removing the angle-dependence for fusion rates would lead to prohibitively high bending energies just after a fusion event. Making the dependence much stricter would cause out-of-equilibrium network fragmentation as fission at a node would often lead to configurations where local fusion is impossible.

Given the role of the angular sensitivity in the fusion rate, increasing *α*_*i*_, *B*_*i*_ in tandem at both degree-2 and degree-3 junctions exponentially decreases the overall association constants at a fixed value of the local fusion prefactor *k*_*ui*_ (Fig. 5A-B). As an alternate approach, we isolate the effect of junction stiffness on the mechanics of the network, varying the bending modulus, *B*_*i*_, while maintaining a fixed *α*_*i*_ in defining the fusion rate. The effect on the association constants is then non-monotonic (Fig. 5C-D). At low fusion rates, the bending stiffness has no effect as the network structure depends on the equilibrium balance between fusion and fission. At intermediate fusion, near the percolation transition, increasing the bending modulus raises the network connectivity (higher *C*_1_, *C*_2_). This effect arises due to the out-of-equilibrium local fusion and fission interactions. High junction stiffness implies that upon fission the edge orientations are very tightly constrained to the optimal values that give the highest local fusion rate. As a result, rapid refusion is very likely and the escape probability is low, giving a notably super-linear increase in association constant as a function of *k*_*ui*_. At the highest fusion rates, when the network is compact and thoroughly interconnected, high bending stiffness constrains the remaining degree-1 and degree-2 nodes on the periphery from being able to reach each other, reducing the overall association constants *C*_*i*_ (Fig. 5C-D).

**FIG. 5.**
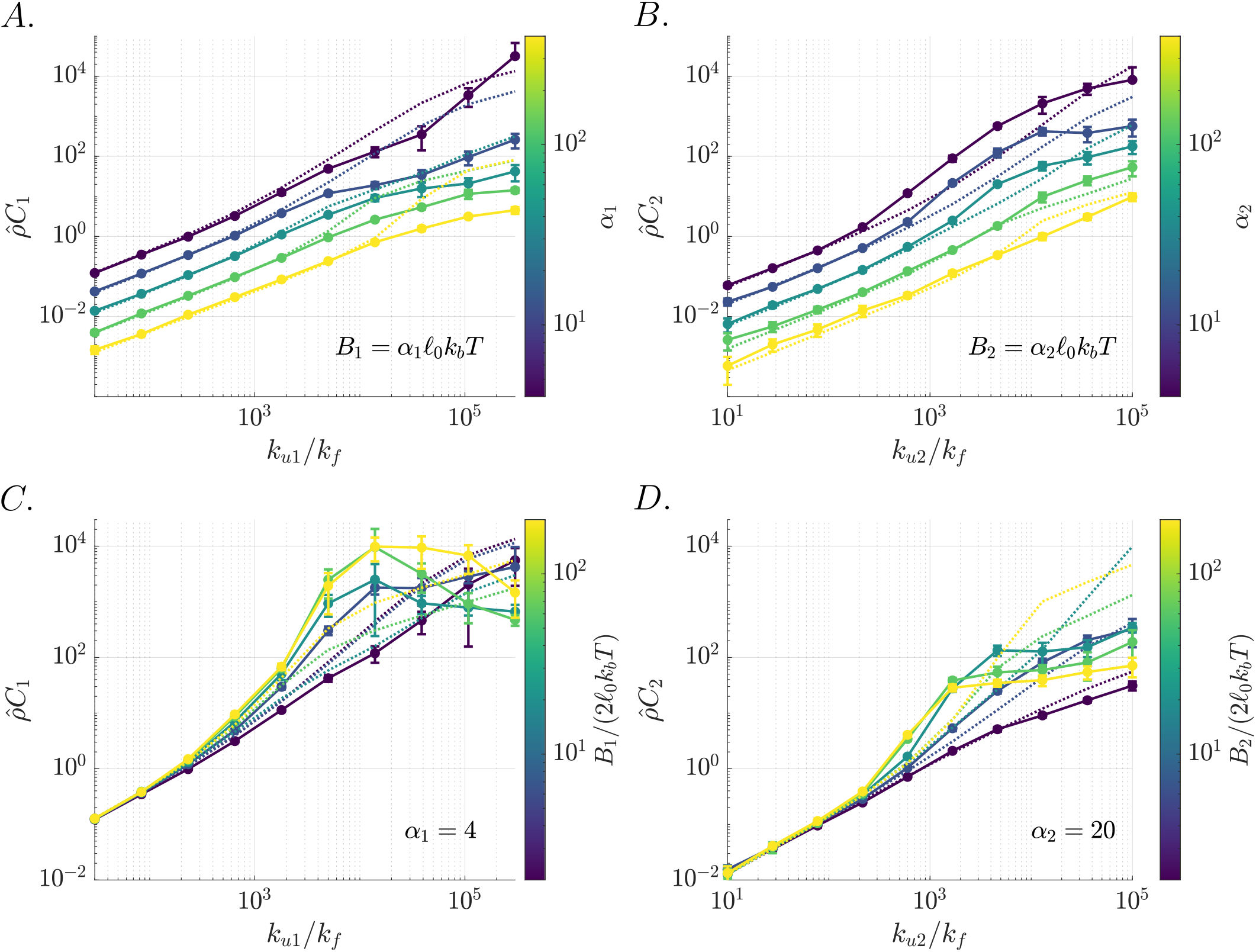
Effect of fusion sensitivity and bending stiffness on network structure. **A**. *C*_1_ is plotted as a function of the local fusion parameter *k*_*u*1_ for different values of *B*_1_, *α*_1_ (colored lines). **B**. *C*_2_ is plotted as a function of the local fusion parameter *k*_*u*2_ for different values of *B*_2_, *α*_2_ (colored lines). **C**. *C*_1_ is plotted as a function of the local fusion parameter *k*_*u*1_ for different values of the bending modulus *B*_1_ (colored lines), while *α*_1_ = 4 is held constant. **D**. *C*_2_ is plotted as a function of the local fusion parameter *k*_*u*2_ for different values of the bending modulus *B*_2_ (colored lines), while *α*_2_ = 20 is held constant. Dotted curves indicate the approximate non-equilibrium association constants estimated from Eq. 7, for each parameter set. All panels use a constant ratio of *k*_*u*2_*/k*_*u*1_ = 1*/*3. Error bars indicate standard deviation across simulation trials.

Overall, increasing the mechanical stiffness of the mitochondrial segments and junctions can reduce network connectivity (by limiting the angles at which fusion can occur) or can enhance it (by increasing the probability of being in a favorable configuration following fission to enable rapid re-fusion). The dominant effect is determined by the microscopic kinetics of how fusion rates depend on the orientations of mitochondrial units.

### E. Mitochondrial motility hinders percolation but enhances maximal network connectivity

We next consider the role of effectively diffusive mitochondrial motion in modulating network architectures. A variety of recent measurements in both eukaryotic and prokaryotic cells have demonstrated that apparent diffusion in the cytoplasm actually relies on active processes within the cell, and is substantially reduced upon ATP depletion [9, 60, 61] or inhibition of various molecular motors [9, 62, 63]. We therefore consider the diffusive motion in our model to be an emergent result of many unspecified spatially distributed active processes.

To investigate the effect of active diffusion on network architecture, we alter the temperature *T* that sets the magnitude of the stochastic Brownian forces 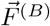, without changing the angle-dependence parameter *α*_*i*_ in the fusion rates. As expected, lowering the temperature at a fixed fusion rate proportionally decreases the effective diffusivity of mitochondrial units (Fig. 6A). Increasing fusion drives down the edge diffusivity, as each mitochondrion becomes more likely to be incorporated into larger network components. The diffusivity drops very low below the network percolation transition, where most edges are incorporated into a single large cluster (see Fig. 2B).

**FIG. 6.**
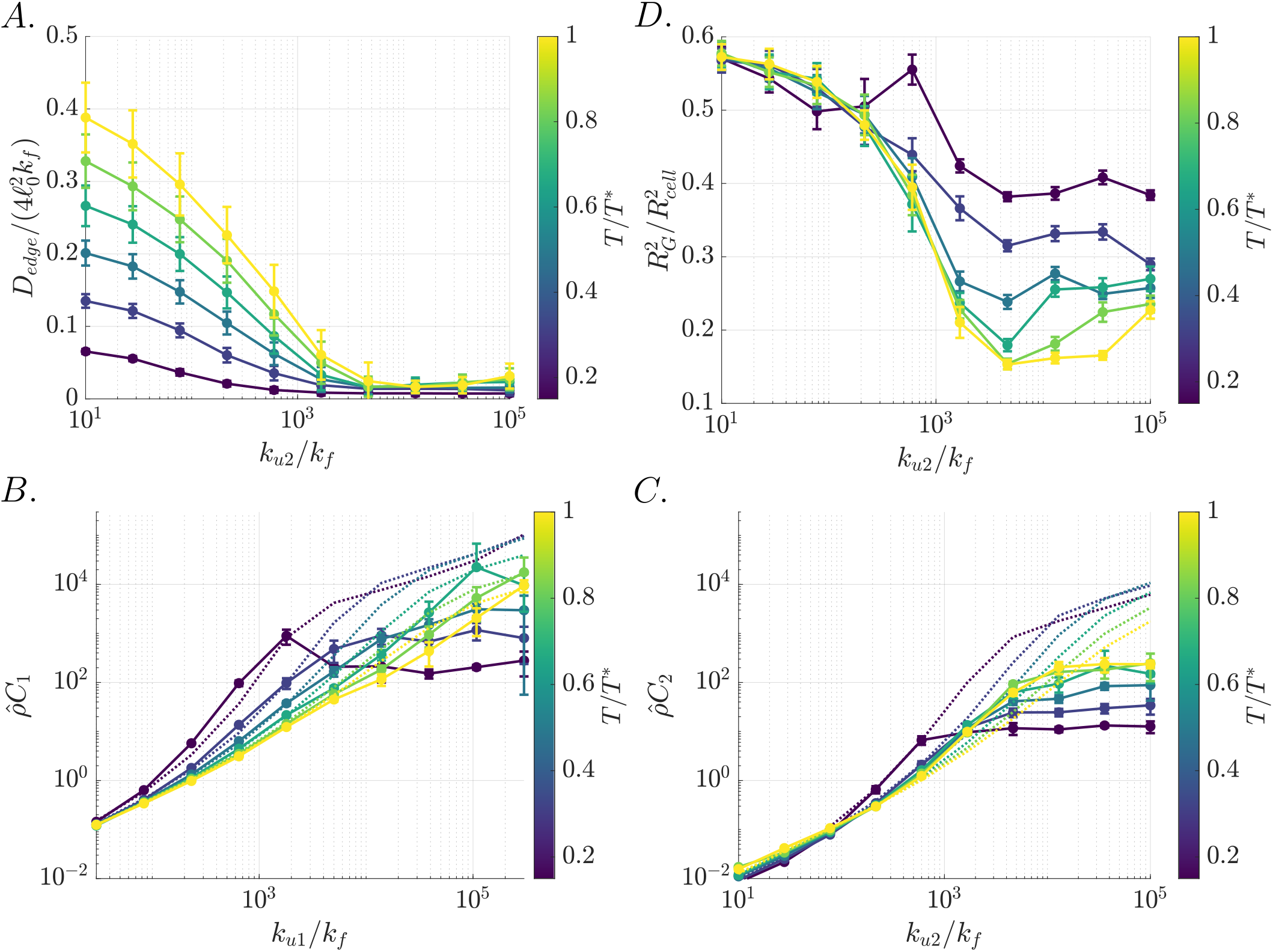
Mobility affects network structure. **A**. The effective edge diffusivity *D*_*edge*_ is plotted as a function of *k*_*u*2_ for different values of the diffusion temperature, *k*_*b*_*T* (colored lines) relative to the default diffusion temperature, denoted *T* ^*∗*^. Highly fused networks result in less mobile mitochondria. **B**. The tip-tip association constant, *C*_1_ is plotted as a function of *k*_*u*1_ for different values of the diffusion temperature, *k*_*b*_*T* (colored lines). **C**. The tip-side association constant, *C*_2_ is plotted as a function of *k*_*u*2_ for different values of the diffusion temperature, *k*_*b*_*T* (colored lines). Dashed lines indicate our (nonequilibrium) analytic predictions. **D**. The radius of gyration *R*_*G*_ is plotted as a function of *k*_*u*2_ for different values of the diffusion temperature, *k*_*b*_*T* (colored lines). Highly fused networks are more compact, but this effect is reduced at lower temperatures, where mitochondria are less mobile and the remodeling process freezes up. A-D use *k*_*u*2_*/k*_*u*1_ = 1*/*3. Error bars indicate standard deviation across simulation trials.

In terms of network structure, decreasing the mitochondrial motion has an analogous effect to increasing the mechanical stiffness of the network. Indeed, lower effective temperature is expected to yield a tighter distribution of bending angles at fused nodes. As a result, in highly percolated networks, lower temperature reduces the large-scale network flexibility and thus limits the ability of peripheral lower-degree nodes to reach each other, reducing association constants (Fig. 6B-C). For intermediate fusion rates, lower temperature results in more favorable edge orientations immediately after fission. This makes re-fusion events more likely to occur before the two nodes can escape from contact, resulting in a higher association constant. This effect is further enhanced by the slower motion of the recently-fissed edges, which also lowers the escape probability. The resulting association constants at low to intermediate fusion rates can be approximately predicted from the simplified model encompassed by Eq. 7 (Fig. 6B-C, dotted lines).

Finally, the magnitude of active diffusion is found to affect the network compactness. At intermediate to high fusion and lower effective temperature, the reduced mobility and flexibility described above lead to increased concentrations of degree-2 nodes, longer segment length, and therefore higher radius of gyration, *R*_*G*_ (Fig. 6D). Higher effective temperature allows for greater flexibility, catalyzing the rearrangements necessary to bring the network to a highly compact state (small *R*_*G*_). Analogous effects have previously been noted for the formation of diffusion-limited aggregates by particles with varying diffusive step sizes [45].

### F. Rapid network rearrangement requires intermediate fusion rates

To gain a sense of the overall rearrangement dynamics of mitochondrial networks, we quantify the rate of tip-tip and tip-side novel fusion in our simulations, defined as the total number of fusion events of each type per unit time that do not involve the connected partner prior to fission. The measured rates of novel fusion show a non-monotonic dependence on the microscopic fusion rates *k*_*ui*_ (Fig. 7A,B). At low *k*_*ui*_ all fusion events are rare, whereas at high *k*_*ui*_ the escape probability following a fission event is low and most fissions thus lead to a rapid re-fusion with the same mitochondrial partner. At high network connectivity, mechanical and spatial constraints prevent distant nodes from reaching each other, so that any fissing pair is very likely to re-fuse rather than seeking out a novel fusion partner. The novel fusion rates peak at intermediate *k*_*ui*_, where contacts between nearby nodes tend to result in successful fusion but escape from the contact volume following fission is still frequent. Interestingly, the extracted microscopic fusion rates for mammalian mitochondrial networks lie reasonably close to the values that yield peak novel fusions. This observation indicates that mammalian mitochondrial networks are not only poised at the boundary of structural percolation, but are also located in a parameter regime that gives high dynamic turnover rates of the network, allowing each mitochondrion to sample different fusion partners relatively rapidly.

**FIG. 7.**
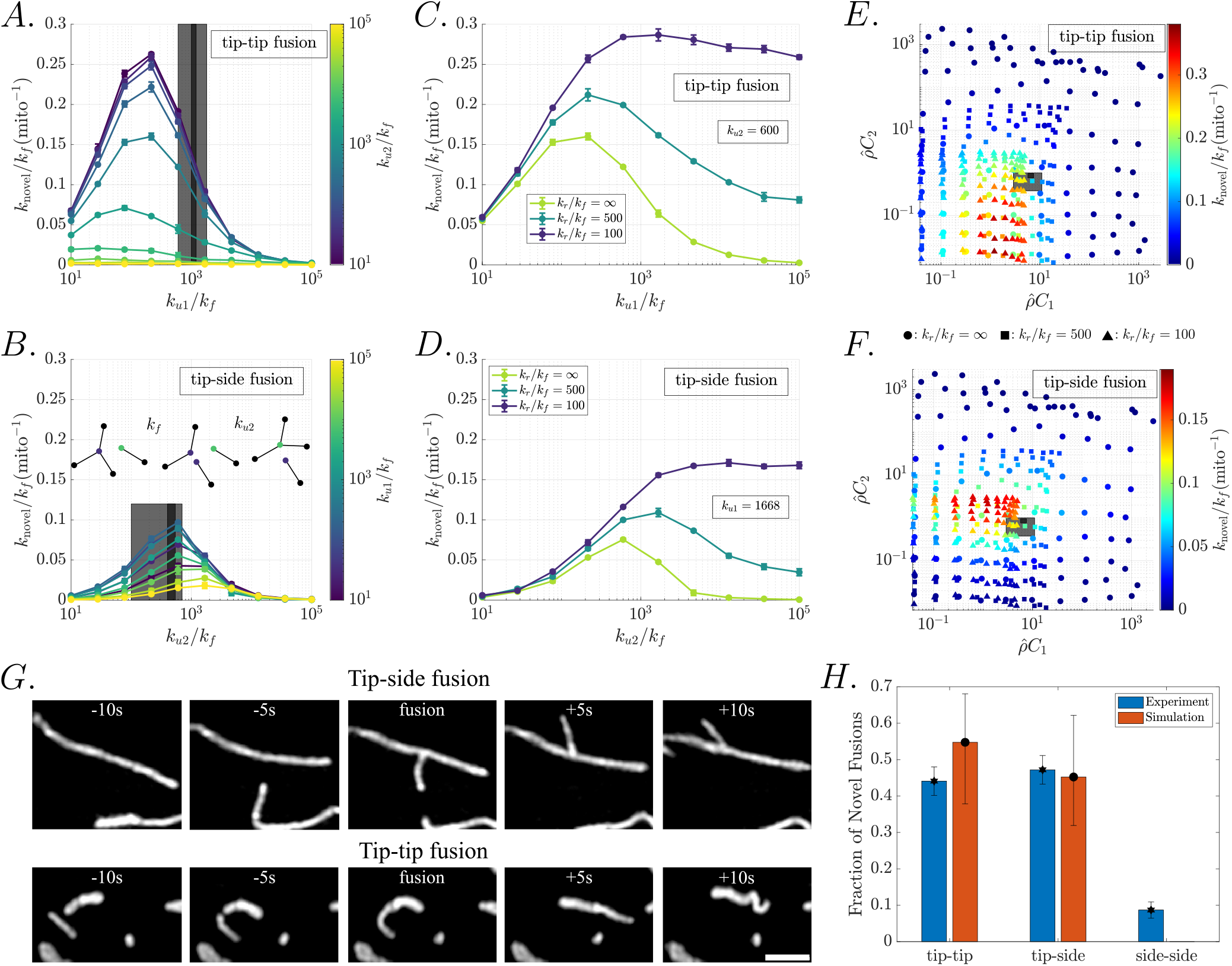
Novel fusions peak at intermediate fusion rates. **A-B**. The rate of novel fusions of each type is plotted as a function of the fusion parameter *k*_*u*1_ (*k*_*u*2_) for different values of the fusion parameter *k*_*u*2_ (*k*_*u*1_) (colored lines). Nodes do not enter an inactive state following fission and may immediately re-fuse. Vertical gray bars indicate the range of *k*_*u*1_(*k*_*u*2_) extracted from experimental networks. Inset: a degree-3 node (purple) undergoes a fission event before tip-side novel fusion with a degree-1 node (green) occurs. **C-D**. The rate of novel fusions of each type is plotted as a function of the fusion parameter *k*_*u*1_(*k*_*u*2_) for different values of the recharge rate, *k*_*r*_ and fixed *k*_*u*2_(*k*_*u*1_) corresponding to the experimentally relevant regime. Nodes enter an inactive state following fission, returning to an active state (in which fusion is allowed) at rate *k*_*r*_. **E-F**. The rate of novel fusions of each type is plotted (colors) as a function of the mean-field association constants, 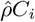 Symbol shape corresponds to different values of the recharge rate, *k*_*r*_. Gray boxes show the experimental values (mean and standard deviation) of 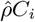 cfor hiPSC [6] (dark gray) and COS-7 cells (light gray). **G**. Examples of observed fusion events between mitochondria in COS-7 cells. Top: tip-side fusion. Bottom: tip-tip fusion. Scale bar: 2*μ*m. **H**. The fraction of fusion events of each type observed in COS-7 cell imaging (blue) and extracted from simulations (red). Error bars: (A-D) standard deviation across trials; (H) for experimental measurements: sampling error for a binomial distribution; for simulations: error bars show range of results corresponding to parameters for the light gray box in E,F.

Given our default simulation time-scale set by *k*_*f*_ = 0.5min^*−*1^, the total rate of novel fusions (using mammalian cell estimates of 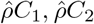, see Fig. 3 gray boxes) is estimated to be 0.06 *−* 0.11min^*−*1^ per mitochondrial unit. These values are comparable with the measured rate of *∼* 4 fusion events per hour per mitochondrion reported for mammalian cells in Ref. [64]. This steady-state novel fusion rate corresponds to a characteristic time for a ‘complete’ fission (a fission event that does not result in re-fusion with the same partner) of *∼* 12min, in agreement with the fission time-scales of 10 *−* 20min used in prior studies [8, 48, 49]. Recently published time-lapse data of mitochondrial network dynamics in hiPSC cells and accompanying network-tracking software [6] yields an estimate of 0.24 *−* 0.32 fusion or fission events per minute per *μ*m of mitochondrial length, also comparable with the simulated values shown here. The matching time-scales for novel fusion justify our selection of the microscopic fission rate constant, and enable the dimensionless results of the model to be interpreted in terms of concrete time units.

Interestingly, imaging of mitochondrial dynamics indicates that mitochondria do not tend to re-fuse immediately following a visible fission event [2, 64]. We note that it is possible that rapid fission and re-fusion cycles do occur but cannot be observed given the limited spatiotemporal resolution of the imaging studies. In this case, the observable fissions and fusions would correspond to the complete fissions and novel fusions of our simulation model. An alternate explanation to the absence of rapid-refusion in imaging studies is that there is an inherent delay before recently fissed mitochondria are able to acquire the machinery necessary for fusion. We therefore consider an extension of the model that prevents immediate refusion by incorporating a “recharge rate”, *k*_*r*_. When nodes undergo fission, they enter an inactive state in which they cannot undergo fusion (i.e. *k*_*ui*_ *→* 0 for each inactive node). Nodes in the inactive state then return to the active, fusion-capable, state with rate *k*_*r*_. The simulations described thus far correspond to instantaneous reactivation, with *k*_*r*_ *→*.*∞*

Slow recharge rates allow newly fissed nodes to escape from their original partner before re-fusing. The separation of two newly-fissed nodes to outside their contact range requires an average of roughly *τ*_esc_ *≈* 0.0024 (see Supplemental Material), indicating that dimensionless recharge rates with *k*_*r*_ ≲ 400 are sufficient to enable nodes to leave the immediate vicinity of their previous fusion partner. Slower recharge rates yield an enhancement in novel fusions when *k*_*ui*_ are high (Fig. 7C,D). However, at low values of microscopic fusion rates *k*_*ui*_, slow recharge can lower the overall rate of fusion (novel and re-fusions both), causing the networks to become less well-connected.

As shown in Fig. 7E-F, the peak novel fusion rates occur at intermediate values of the association constants. Although slower recharge boosts the peak rate of novel fusions, the overall dependence on the association constants is the same for a broad range of recharge rates. Specifically, the maximum rate of novel fusion occurs when 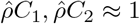, corresponding to a system where the fusion and fission are closely balanced. The experimentally extracted association constants for mammalian networks are 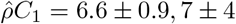 and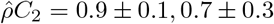, which gives novel fusion rates that are approximately 60% of the maximum possible values. Notably, these values of 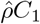 cannot be reached with a recharge rate of *k*_*r*_ = 100, setting a lower limit on the rate at which mitochondria are able to recover and engage in subsequent fusion events.

The predicted rates of tip-tip versus tip-side novel fusions are comparable (0.05 *±* 0.02, 0.04 *±*0.01mito^*−*1^min^*−*1^, respectively, based on measured association constants for COS-7 cells), as previously reported for mitochondrial dynamics in mammalian cells [64]. To further test this prediction, we perform an analysis of fusion events via live imaging of COS-7 cells, labeling each event as either tip-tip, tip-side, or (rarely) side-side (see Fig. 7G). We find that roughly 44% of 161 observed fusions are tip-tip, while 47% are tip-side, in good agreement with the extracted simulation-based novel fusion rates (Fig. 7H). This near-equivalence of novel fusions for the two event types is observed even though the microscopic fusion rate constants (*k*_*u*1_, *k*_*u*2_) differ by a factor of 3. The distinction arises because the novel fusion rate incorporates geometric and mechanical constraints that affect the distribution of angle-dependent fusion rates as well as the ability of newly-fissed nodes to escape from each other.

Overall, our results show that mammalian mitochondrial networks lie in a parameter regime that enables them to be highly ‘social’ [65, 66], rapidly encountering novel partners for both tip-tip and tip-side fusions.

## IV. CONCLUSIONS

In this work, we present a framework for understanding how cell-scale mitochondrial network structure arises from the dynamics of individual organelle units. Mitochondrial networks vary widely, from ‘social’ networks of interacting but physically fragmented units [65, 66] to highly connected tubular networks [10, 28, 67], depending on cell type, genetic perturbations, and environmental conditions. The spatially-resolved model presented here elucidates the impact of underlying microscopic rate constants and mechanical features on the morphology of these networks.

The network architecture is shown to be highly dependent on the ratio of microscopic fusion and fission rates for tip-tip and tip-side interactions. Analysis of published data for both mammalian and yeast cell networks indicates that the two microscopic fusion rates are relatively well-balanced (*k*_*u*2_*/k*_*u*1_ *≈* 0.3), resulting in structures with modest segment lengths. While mammalian networks are poised at the edge of the percolation connectivity transition, yeast networks are generally highly connected structures. We demonstrate that this distinction arises from the geometric confinement and effectively high density of mitochondria at the surface of the yeast cell, without requiring substantially different microscopic rate constants.

Using a mass-action model, we show that the effective mean-field association constants (*C*_1_, *C*_2_) that describe network architecture [28, 35] can be analytically predicted from microscopic parameters and geometric constraints. Crucially, the standard mass-action model breaks down at intermediate to high fusion rates, where spatial transport becomes limiting. At intermediate rates there is an enhancement in the probability of rapid re-fusion following each fission event, due to the fact that recently fissed edges are pre-aligned to facilitate fusion. This is a non-equilibrium feature, resulting in a superlinear dependence of association rates on microscopic fusion to fission ratios. The model thus serves as an example of the general principle that an input of energy into a biomolecular system can amplify specific states [68] and sharpen the system’s response to small parameter changes [69, 70].

At high fusion, the formation of spatially extended structures precludes the ability of dangling ends to find each other, resulting in a plateau of the association constants. The mechanical flexibility of mitochondrial junctions, as well as the motility of mitochondrial units, both suppress re-fusion rates at intermediate *k*_*ui*_ and increase the maximal network connectivity at high *k*_*ui*_.

We further quantify the rate of novel fusions, wherein a mitochondrial unit switches to another fusion partner, rather than re-fusing rapidly with the same partner after a fission event. Such novel fusion rates provide a metric for network rearrangement time-scales, and are maximized at intermediate values of the microscopic fusion, somewhat below the percolation transition. Mammalian mitochondrial networks are shown to operate in a parameter regime with relatively high rates of novel fusion, allowing for constant rearrangement of network structures. Thus, mammalian mitochondria can be thought of as highly ‘social’ organelles [65, 66], with frequent interactions and exchanges between distinct segments.

We note that in experimental studies mitochondria are generally not observed to fuse again immediately after fission [64]. However, the spatiotemporal resolution of 3D imaging is inherently limited (usually with frame rates below 10 Hz), so that rapid fission and re-fusion events may still occur without being directly observable. Such rapid refusion could be supported by the convergence of fusion and fission machinery at ER-mitochondrial contact sites, enabling fast successive dynamics [71]. In this case, the ‘novel fusion’ and corresponding ‘complete fission’ rates predicted by our simulations (rather than the microscopic fusion and fission rate constants) are more directly comparable to experimental measurements. Alternately, there may be additional transitions that must occur before a newly fissed mitochondrion is capable of fusing again, explicitly precluding rapid re-fusion events. We thus incorporate into our model a latent recovery period, which enhances the rate of novel fusions at high values of *k*_*ui*_, but also limits the overall association constants and hence the connectivity of the network. Regardless of the microscopic fusion and recovery kinetics, the rate of novel fusion is shown to be maximized at intermediate association constants 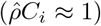, where overall fusion and fission rates are approximately balanced. Further, the association constants obtained for mammalian cells are shown to correspond to approximately equal rates of novel tip-tip and tip-side fusions, as observed in live-cell imaging measurements.

This work takes a key step towards gaining a fundamental understanding of dynamic spatial networks because it forms a direct link between observable macrostates and tunable microscopic parameters, while accounting for spatial and mechanical constraints. Experimental perturbations that are known to alter global mitochondrial dynamics [10, 20, 28] may do so through a combination of modulating fusion and fission rate constants, mitochondrial flexibility, and transport. The modeling results presented here make it possible to link the observed cellular-scale changes in network structure with experimentally accessible mechanistic factors.

The model described here sets the stage for addressing a number of open questions concerning dynamics on and of the mitochondrial network. First, how does network morphology control and affect the distribution and dispersion of proteins, nucleoids, ions, and lipids within the mitochondrial compartments? The spatiotemporally resolved simulations form a natural platform for tracking the spread of intra-mitochondrial material. Second, how efficiently can mitochondria be sorted, during cell division or through quality control mechanisms, based on their connectivity and material contents? Sorting of healthy mitochondria into daughter buds has been proposed as a mechanism contributing to the aging of budding yeast cells [72]. Selective destruction of unhealthy mitochondria excluded from the network structure is also thought to be a major component of mitochondrial homeostasis [2, 73] and could be incorporated into the model. Third, the spatial architecture of the mitochondrial network may modulate the distribution of interactions with other cellular structures, including the localization of ribosomes to the mitochondrial surface [74] and the formation of contact sites between mitochondria and the endoplasmic reticulum [75]. Understanding the connectivity and transport of mitochondria could elucidate how such contacts are maintained and how they respond to changing conditions. Overall, our simulations provide an *in silico* framework for exploring the interplay of structure and dynamics in the mitochondrial population, furthering our understanding of how the observed morphology of these organelles couples to their many biological functions.

## V. METHODS

### A. Mechanical and dynamic model for the mitochondrial network

#### 1. Mitochondrial mechanics

Individual mitochondrial units are represented as edges of ground-state length *l*_0_ = 0.5*μ*m and steric radius *r*_*s*_ = 0.1*μ*m. When the units are connected into larger structures, nodes of degree 2 and 3 mark the connection points. Whenever an edge terminates at a degree 1 node, the node is placed at an inset distance of *r*_*s*_ along the length of the edge. The length of the edge (*l*) is then defined to extend to the tip of the surrounding steric spherocylinder, so that the ground-state value of this length remains at *l*_0_ regardless of the edge connectivity.

The time-evolution of the nodes is governed by an overdamped Langevin equation (Eq. 1), incorporating mechanical forces associated with bending, stretching, steric interaction, and confinement energies for the edge configurations. The mechanical model is analogous to that commonly used for simulating bead-rod chains at the macromolecular scale [76, 77]. The stretching energy for each individual unit is set to

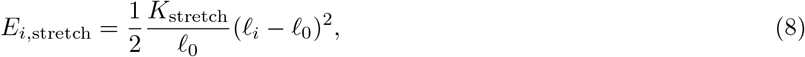

where *l*_*i*_ is the length of edge *i*, and *K*_spring_ is the stretching spring constant, set sufficiently high that the root mean squared (RMS) deviation in edge length is below 0.05*l*_0_.

The bending energy at each junction node is given by Eq. 3 and depends on the angles between edges connected at that junction. For a degree-2 node, *θ*_*i*_ is the angle between the two edges and the bending energy is lowest at *θ*_*i*_ = 0. For a degree-3 junction, the three angles 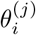 between the adjacent edges minimize the bending energy in a symmetric planar configuration with 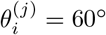. The bending moduli *B*_1_, *B*_2_ scale the energetic penalty for deviating away from the ground-state angles, and their default magnitude is estimated from experimental data as described in Supplemental Material.

A steric energy *E*_*i,j*,steric_ is defined for each pair of edges *i, j* that are not directly connected to each other. We compute the minimal distance *d*_*i,j*_ between the line segment connecting the nodes of edge *i* and the line segment connecting the nodes of edge *j*. The steric energy is set to give a quadratic penalty when this minimal distance is less than twice the steric radius:

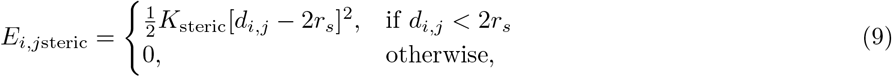

where the steric modulus *K*_steric_ is set sufficiently high that the RMS value of |*d*_*i,j*_ *−* 2*r*_*s*_| for sterically interacting edges is less than 0.02*l*_0_. This form of the steric energy effectively treats individual fragmented units as spherocylinders of radius *r*_*s*_ and total length (to the tips of the spherical caps) equal to *l*_0_. Edges connected into longer tubes overlap their hemispherical end-caps so that the non-connected mitochondria are excluded from a continuous region of radius *r*_*s*_ surrounding the entire tube.

The final mechanical component of the model is the confinement force. Simulations are carried out in a spherical domain of radius *R*_cell_ = 10*l*_0_, with a steep quadratic energy penalty for mitochondrial nodes that step outside the domain. For surface-bound simulations representing yeast mitochondria, we use *R*_cell_ = 4.6*R*_0_ and an additional hookean confinement force provides a tethering of the mitochondria to the inside surface of the domain. The confinement energy is thus defined for each node *i*:

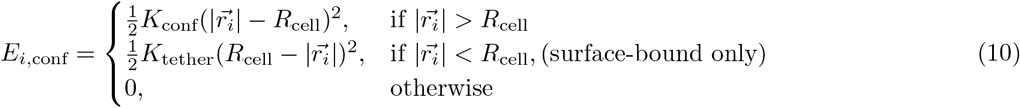

All 3D mitochondrial network simulations include 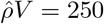 edge-units in the spherical domain. All surface-bound simulations include 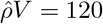 edge-units, giving a similar volume density to that found in mammalian cells and yeast cells, respectively.

#### 2. Network dynamics

At each step of the simulation, the total mechanical energy for all nodes and edges is computed, and the mechanical force 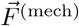 is found as the spatial derivative of that energy with respect to node positions (Eq. 2). In addition, the Brownian force is computed as 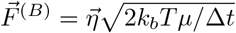 with Δ*t* the simulation timetep, *μ* the friction coefficient of each node, *k*_*b*_*T* the effective thermal energy, and each dimension of 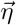 a normally distributed random variable with zero mean and unit variance. We non-dimensionalize length-units by 2*l*_0_, time units by 1*/k*_*f*_, and force units by *μ*. The default effective thermal energy is set to *k*_*b*_*T* = 1 in these dimensionless units. The simulations are integrated forward using a fourth-order Runge-Kutta method [78] (previously validated for Brownian dynamics simulations [79]) to update the node positions with timestep Δ*t* = 10^*−*4^.

At each time-step of the simulation, a pair of nodes is able to undergo fusion if they are within contact distance 2*r*_*c*_, where *r*_*c*_ = 0.15 is taken as the contact radius. The edge-units they belong to must not be connected and should be well-aligned. The rate constants for fusion depend on the bending angles that would arise at the newly formed junctions, as given in Eq. 4. The probability of a pair fusing during time step Δ*t* is computed according to:

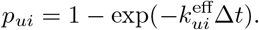

Similarly, during each time-step, each degree-2 and degree-3 node have a probability of fission given by

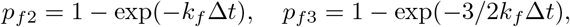

respectively. Upon fission, the newly formed nodes are placed just outside of steric overlap (*d*_*i,j*_ = 2*r*_*s*_). Details of node placement upon fusion and fission, as well as angle calculations defining the fusion rates, are given in Supplemental Material. At most one fission or fusion event can occur per each pair of nodes within a single timestep.

Simulations were initialized with a fully fragmented network and were run for at least 1.5 *×* 10^6^ timesteps. The last 30% of the time-course was used to compute steady-state metrics, and the results were averaged over at least 3 independent trials for each simulation.

### B. Cell culture, imaging, mitochondrial network extraction, and fusion quantification for COS-7 cells

#### 1. Network Extraction

COS7 cells were grown in Dulbeccos modified Eagles medium (DMEM) supplemented with 10% fetal bovine serum and maintained in culture for a maximum of 20 passages. 72h before imaging, cells were seeded in a 6-well plate at 70,000 cells per well. 48h before imaging, cells were transfected with mito-mScarlet (pMTS mScarlet N1), a gift from Dorus Gadella (Addgene plasmid #85057) with Lipofectamine 2000 (Invitrogen) [80] in Opti-MEM medium (Invitrogen) following the manufacturers instructions. Cells were grown and handled as in [20].

To facilitate imaging of whole mitochondrial networks, cells were seeded on fibronectin micropatterned coverslips (made as described here [81]) the evening before imaging (approximately 16h).

Mitochondrial networks were acquired on a Nikon Eclipse Ti2-E inverted microscope spinning disk CSU-W1 equipped with an sCMOS camera (Photometrics Prime 95B) and a 100x 1.49 NA oil immersion objective (SR HP Apo TIRF 100x/1.49, Nikon). Images were acquired in 100nm z-steps at 37C and 5% CO2 in phenol-red-free DMEM. Before further processing, the RAW images were cropped and deconvolved with Huygens Core, offered by the EPFL imaging core facility (BIOP), using a theoretical PSF.

To receive the graph output the deconvolved z-stacks were processed by Mitograph [82] running via the Windows subsystem for Linux (WSL2). Tiff stacks were processed with the following command:./MitoGraph -xy 0.11 -z 0.1-labels off -path *<*file-path*>*.

#### 2. Fusion Quantification

The following plasmids and reagents were used to label mitochondria in COS-7 cells. Mito-GFP (Cox-8 presequence) was a gift from H. Shroff (NIH, Bethesda, USA), FASTKD2-EGPF was a gift from J.-C. Martinou, mtAT1.03 was a gift from Hiromi Imamura (Kyoto University, Japan), mito-TagRFP was generated in the laboratory from mito-GFP, TOM20-mCherry, TOM20-EGFP, and TMRE (Sigma, 87917, 500nM for 10min and rinsed by PBS). In addition to fluorescent labelling, mitochondria were also imaged by phase-contrast. Transfections were performed using Lipofectamine 2000 (Invitrogen, 11668027) following the manufacturers protocol. Live-cell time series were acquired on a custom-built instant structured illumination microscope (iSIM) [83], a 3D NSIM Nikon inverted fluorescence microscope (Eclipse Ti), a Leica TCS SP8 inverted confocal microscope and a Zeiss Axio Observer 7 inverted widefield microscope. Fusions were manually detected and classified by fusion type using Fiji [84]. Mitochondrial fusions were defined as events where two individual mitochondria came together and formed a single unit that remained clearly connected for several frames following the fusion event. For visualization, noise was removed from representative images with a 1-pixel mean filter. Some data used for this fusion quantification was reused from Rey et al. [85] and Kleele et al. [20].

### C. Quantifying experimental network structures

The three data sets referenced in this work are: 1. A set of 93 consecutive network structures for a single hiPSC cell (3.253 seconds per frame), published in Ref. [6] and available on the MitoTNT github page. 2. A set of 2878 *S*.*cerevisiae* budding yeast networks, of which 2794 are used for the full analysis (keeping only networks with *ρ*_1_, *ρ*_2_, *ρ*_3_ *>* 0). These structures are accessible via the Mendeley Data webpage associated with Ref [10]. 3. A set of 37 structures from COS-7 cells, obtained as described above. Custom Matlab data structures for handling and visualizing networks are provided at https://github.com/lenafabr/networktools.

In order to compare our simulated networks to experimental data, we compute an approximate mitochondrial volume density for both mammalian-cell data sets. The total mitochondrial length is found as the sum of the edge lengths in each network. For the hiPSC and COS-7 mammalian networks, we found mean mitochondrial lengths (plus/minus standard deviation): *L* = 360 *±* 10*μ*m and *L* = 700 *±* 200*μ*m, respectively. As the precise cytoplasmic volumes accessible to the mitochondria were not directly available from the experimental data, we used a convex hull approach (MATLAB’s “boundary” function) to approximate the cytoplasmic volumes. This gave mean volumes: 1500*±*100*μ*m^3^ and 3000 *±* 1000*μ*m^3^, respectively. The corresponding length density of mitochondria is then 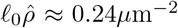. We use a similar density in our simulations, where 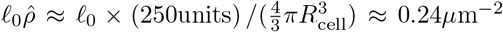. For yeast cells, the distribution of mitochondrial lengths was double-peaked, and our density estimates were taken from the subset of cells corresponding to the peak with larger mitochondrial length. For these cells, the average mitochondrial length was *L* = 60 *±* 20*μ*m (mean *±* standard deviation) As yeast mitochondria reside predominantly on the inner surface of the cell membrane, we used cell surface area (provided in the dataset) to set the size of our simulated cells. The average surface area for the analyzed cells was 70 *±* 20*μ*m^2^, corresponding to a simulation sphere of radius *R*_cell_ *≈* 2.3*μ*m.

To calculate the mean segment length ⟨*L*_seg_ ⟩and largest cluster size fraction ⟨*L*_*G*_*/L*⟩ for the imaged datasets, we use the same procedure as outlined for simulated networks. Estimating association constants *C*_*i*_ for experimental networks requires first computing the equivalent number of nodes in the imaged network structure. The mitograph structures are analyzed to count the number of degree-1 nodes (*X*_1_) and the number of higher-degree junctions (*X*_*k*_ for *k ≥* 3), as well as the total length *L* of the mitochondria. We then compute the effective number of mitochondrial units (*N* ^eff^) and effective degree-2, degree-3 nodes 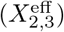 as follows:

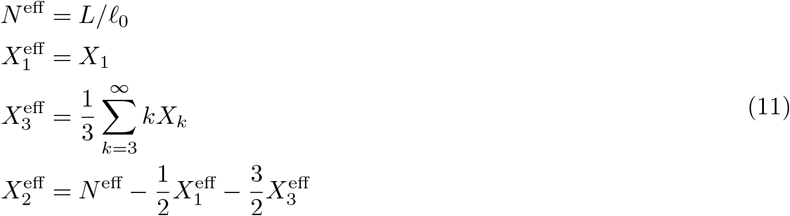

This conversion enables us to calculate 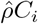 for each experimental network in a way that accurately reflects the connectivity of the mitochondria.

### D. Estimation of contact volume and mean fusion rate for equilibrium association constants C_i_

#### 1. 3D networks

The extent of fusion into degree-2 and degree-3 junctions can be described by the effective association constants *C*_1_, *C*_2_, defined by the steady-state ratio of global fusion and fission events (Eq. 6) or equivalently by ratios of nodes of different degrees. The global fusion rate in a well-mixed equilibrium system should be determined by the microscopic angle-depedent fusion rates 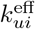, and the volume of the contact zone within which fusion is possible.

The contact volume *v*_cont,i_ is the volume surrounding a degree-1 (or 2) node in which a degree-1 node could be placed such that a valid tip-tip (or tip-side) fusion could occur. This volume is limited by a sphere of radius 2*r*_c_, which defines the contact distance between two nodes. We take *r*_c_ = 0.15*μ*m throughout. The steric exclusion between two mitochondrial segments reduces the actual configurational volume within which two nodes can fuse. For tip-tip fusion, this reduction involves subtracting the volume of a spherocylinder tip of radius 2*r*_s_ (where *r*_*s*_ = 0.1*μ*m). For tip-side fusion we consider the approximation of a perfectly straight existing degree-2 junction to subtract a cylindrical volume of radius 2*r*_s_. The contact volumes for each type of fusion are then given by:

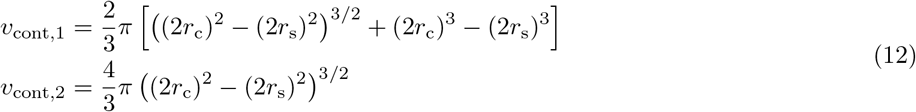

The quantity, 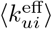, represents the mean fusion rate for a pair of nodes, given that the pair is within contact range. This rate depends on the orientations of the edges attached to the prospective fusing nodes. For tip-tip fusion, we consider an existing edge aligned along the z axis with a second edge fusing in at angle *θ*. The fusion rate 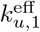 (Eq. 4a) is averaged over all incoming edge orientations, assumed uniformly distributed, to give:

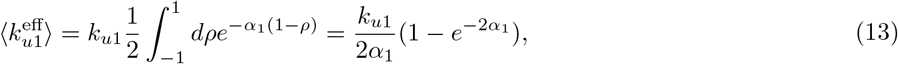

where *ρ* = cos *θ* and *α*_1_ is the angular sensitivity parameter. The resulting equilibrium association constant 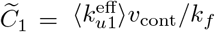 defines the linear estimate plotted in Fig. 3A. The correction factors *p*_esc_, 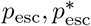 accounting for non-equilibrium effects are described in Supplemental Material.

For tip-side fusion, we employ an analogous strategy to estimate the effective fusion rate. The degree-2 node is placed at the origin and the incoming degree-1 node is taken to be at distance 2*r*_*s*_ + *δ* (along the negative x-axis), where *δ* is set to the average value for a uniform distribution within the contact volume. The first edge is taken to lie along the z-axis, and the second edge to be connected at angle *θ*^(1)^, in the xz plane. The incoming third edge is taken to have polar angle *θ*_2_ and azimuthal angle *ϕ* relative to the plane formed by the first two edges. To avoid steric overlap, the azimuthal angle is limited to be between (*ϕ*^*∗*^, 2*π − ϕ*^*∗*^) where sin *ϕ*^*∗*^ = 2*r*_*s*_*/*(2*r*_*s*_ + *δ*). Junction configurations are not equally likely and must be weighed by the appropriate Boltzmann factor depending on the bending angle *θ*^(1)^ and bending stiffness *B*_1_. We therefore compute:

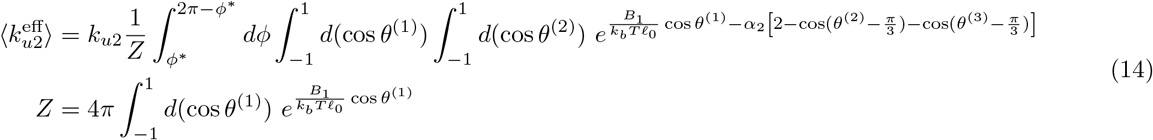

where *θ*^(1)^ is the angle between the two connected edges, and *θ*^(2)^, *θ*^(3)^ are the angles of the incoming edge with each of the two connected ones. The resulting equilibrium association constant 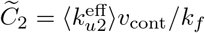 defines the linear estimate plotted in Fig. 3B.

#### 2. Surface-bound networks

For surface-bound networks representing yeast mitochondria, the contact volume becomes an area, and the mean fusion rate is averaged over two-dimensional configurations. For tip-tip fusion, the contact area corresponds to a circle of radius 2*r*_c_ minus an excluded region equal to a rectangle of width 4*r*_*s*_ and a half-circle of radius 2*r*_*s*_ For tip-side fusion, we again take the limit of a perfectly-straight existing degree-2 junction and subtract a rectangle of width 4*r*_s_ from a circle of radius 2*r*_c_. The contact areas are then given by the following expressions:

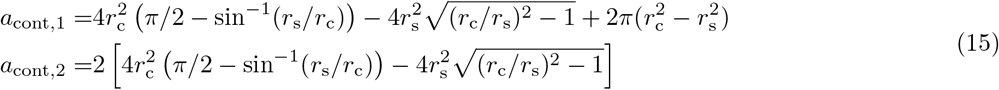

Due to the higher segment density in the surface-bound case, we explicitly subtract from the total available area the space that is sterically excluded by existing nodes. Specifically, we compute 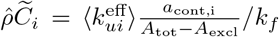 where 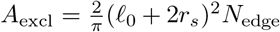 is the approximate excluded area of two rectangles (see [86] Fig.14.17) and 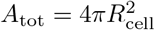. In the 3D case, the corresponding correction was found negligible and was ommitted for simplicity.

We also exclude the sterically inaccessible orientational states when integrating over possible incoming node positions and edge orientations to find the average fusion rate. For tip-tip fusion, we take the node separation to be 2*r*_*s*_ + *δ* as in the 3D calculation above. The first edge lies along the negative x-axis with one node at the origin. Equal weights are assigned to all positions 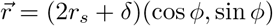 and orientations, *θ*:

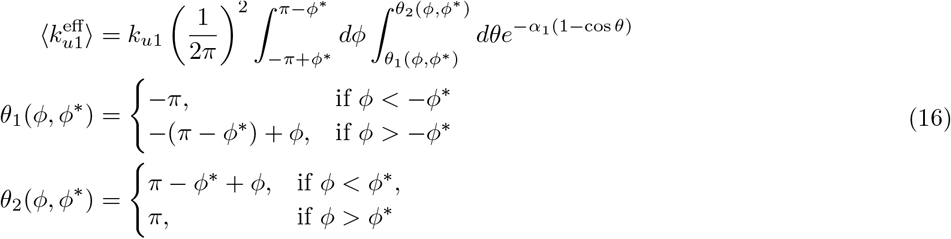

where sin *ϕ*^*∗*^ = 2*r*_*s*_*/*(2*r*_*s*_ + *δ*) as defined above and integration limits are chosen so as to eliminate sterically forbidden node positions and edge orientations. This result corresponds to the linear prediction in Fig. 4C.

For tip-side fusion, the effective average fusion rate within the contact volume is estimated as follows. We place the incoming degree-1 node at a distance 2*r*_*s*_ + *δ* along the positive y-axis, with the degree-2 junction at the origin. The first edge lies along the negative x-axis, with the second edge connected at angle *θ*^(1)^ The angles *θ*^(2)^, *θ*^(3)^ indicate the orientation of the incoming 3rd edge relative to the first and second edges, respectively. Sterically forbidden orientations are approximately eliminated via the integration limits:

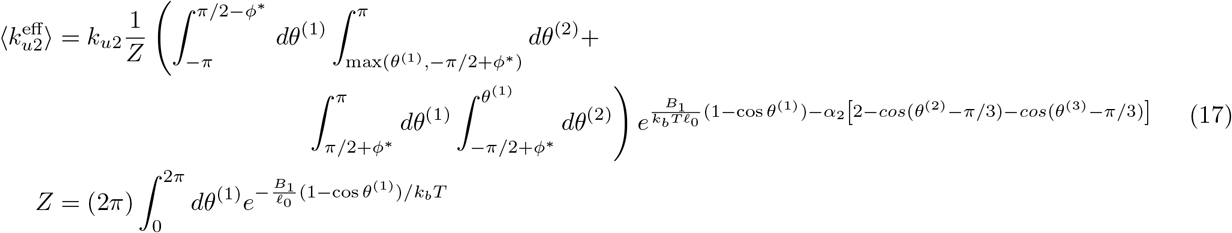

This result gives the linear prediction shown in Fig. 4D.

### E. Solution to the simplified kinetic model for escape

Here we present the solution to the model diagrammed in Fig. 3B (inset), which describes the possible states of a single pair of nodes capable of undergoing fusion and fission. The steady-state kinetic equations for this system are:

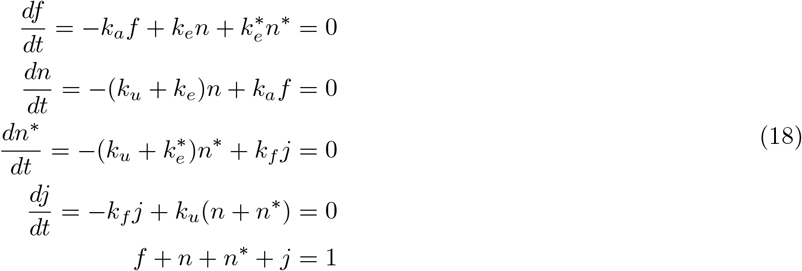

where the variables *j, n, n*^*∗*^, *f* are the probabilites of finding the pair in the corresponding state. The association constants *C*_1_, *C*_2_ (scaled by the volume of the domain) measure the ratio of fused to unfused node pairs, and may therefore be written generically in terms of the state probabilites:

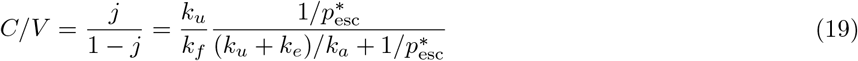

Where 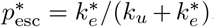. A simpler equilibrium system obeying detailed balance would be one where fission returns the node pair to the same state it was in upon fusion, so that *n*^*∗*^ *→ n* and 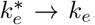 In this equilibrium case, the fraction of disjoint pairs that are close enough to enable fusion must equal the volume fraction of the contact zone: *n*_*eq*_*/*(*f*_*eq*_ + *n*_*eq*_) = *k*_*a*_*/*(*k*_*a*_ + *k*_*e*_) = *v*_cont_*/V*. The association constants may then be expressed as:

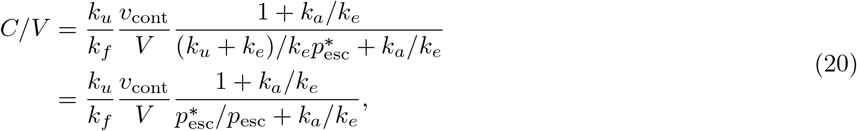

where *p*_esc_ = *k*_*e*_*/*(*k*_*u*_ + *k*_*e*_) is the escape probability for the equilibrated reaction which retains no memory of prior fission.

In the relevant limit of the domain size much larger than the size of an individual mitochondrion, we have *v*_cont_ *« V* and hence *k*_*a*_ *« k*_*e*_. Consequently, we can estimate the overall association constants as:

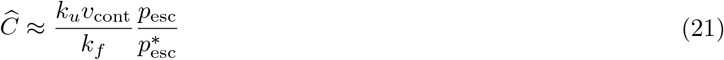

We note that escape is less likely immediately after a fission event, because the nodes are more closely positioned (separation 2*r*_*s*_) than they were upon fusion and the orientation of the segments tends to be more favorable for refusion. Because the correction factor 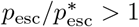 itself increases with rising fusion rates, it allows for a super-linear dependence of the associaton constant on the microscopic fusion rate constant *k*_*u*_.

In this simplified model, the fusion rate *k*_*u*_ incorporates rates 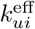 averaged over possible orientations of newly approaching segments (Eq. 13, 14, 16, 17). The escape probabilities must be computed by considering the initial separation of the two nodes within contact range, and their alignment for subsequent refusion. Approximations for these quantities are derived from a reaction-diffusion model in spherical coordinates, as detailed in Supplemental Material.

## IV. ACKNOWLEDGEMENTS

We thank Zubenelgenubi Scott, Padmini Rangamani and members of the Rangamani group for helpful discussions. This work was supported by NSF grant #2310229 and an award from the Chan Zuckerberg Initiative Foundation.

