## Supplemental Material for "Spatiotemporal Modeling of Mitochondrial Network Architecture"

### SUPPLEMENTARY MATERIAL:
### Spatiotemporal Modeling of Mitochondrial Network Architecture

#### S1. IMPLEMENTATION OF FUSION AND FISSION EVENTS

This section details the precise positioning of nodes defining the mitochondrial edge units after a fusion or fission event. The approach described here precludes extremely high steric forces immediately after fission. It also ensures that a very rapid fusion and fission cycle (without intervening mechanical relaxation) would result in little net change in the node positions.

##### S1A. Tip-tip fusion and degree-2 fission

Suppose two edges $i$ and $j$ connect nodes at positions $\left\{\vec{r}_i^{\,(1)}, \vec{r}_i^{\,(2)}\right\}$ and $\left\{\vec{r}_j^{\,(1)}, \vec{r}_j^{\,(2)}\right\}$, respectively, with the first nodes being of degree 1. The two edges can undergo tip-tip fusion when $\vec{r}_{i,j}^{\,(1)}$ approach within contact range $2r_c$, where $r_c = 0.15\mu$m is the contact radius. The newly-fused node $\vec{r}^*$ is set to be halfway between the effective end-points (Fig. S1A). The rate constant $k_{u1}^{\text{eff}}(\theta)$ for the fusion event (Eq. 4a) depends on the angle $\theta$ defined around the putative newly formed node

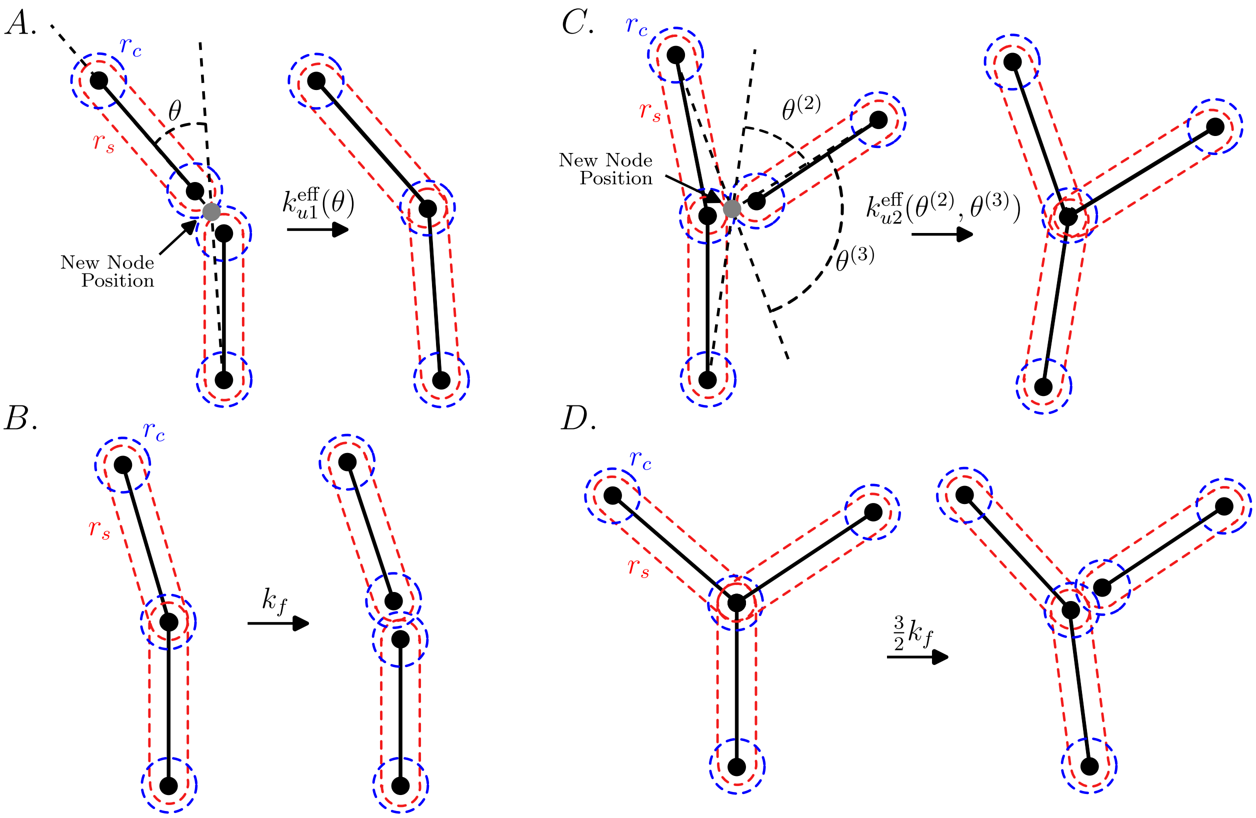


FIG. S1. Details of fusion and fission events as implemented in our model. **A.** Tip-tip fusion. **B.** Degree-2 fission. **C.** Tip-side fusion. **D.** Degree-3 fission.

according to:

$$\vec{r}^{\,*} = \left(\vec{r}_i^{\,(1)} + \vec{r}_j^{\,(1)}\right)/2$$
$$\cos\theta = \frac{\left(\vec{r}^{\,*} - \vec{r}_i^{\,(2)}\right)\cdot\left(\vec{r}_j^{\,(2)} - \vec{r}^{\,*}\right)}{\left|\vec{r}^{\,*} - \vec{r}_i^{\,(2)}\right|\left|\vec{r}_j^{\,(2)} - \vec{r}^{\,*}\right|} \tag{S1}$$

Upon fission of a degree-2 node, the newly formed degree-1 nodes are both placed a distance of $r_s$ away from the position of the original fissing node, while maintaining the same edge orientation (Fig. S1B). If the original connected edges were linearly aligned ($\theta = 0$), this placement results in no steric overlap between the newly separated edges. In the case where they are bent at an angle, a small amount of steric overlap occurs. Because the degree-2 nodes have a substantial bending stiffness, consecutive connected units remain relatively well-aligned throughout the simulation and the steric forces following fission are small.

##### S1B. Tip-side fusion and degree-3 fission

The fusion of a degree-1 node and a degree-2 node also requires the two nodes to be within distance $2r_c$. Suppose $\vec{r}_i^{\,(1)}$ is the position of the degree-1 node on edge $i$ and $\vec{r}_j^{\,(1)} = \vec{r}_k^{\,(1)}$ is the position of the degree-2 node connected to edges $j, k$. The newly formed degree-3 node is again placed at $\vec{r}^*$ halfway between them (Fig. S1C), and the relevant angles determining the fusion rate (Eq. 4b) are defined as follows:

$$\cos(\pi - \theta^{(2)}) = \frac{\left(\vec{r}_i^{\,(2)} - \vec{r}^{\,*}\right)\cdot\left(\vec{r}_j^{\,(2)} - \vec{r}^{\,*}\right)}{\left|\vec{r}_i^{\,(2)} - \vec{r}^{\,*}\right|\left|\vec{r}_j^{\,(2)} - \vec{r}^{\,*}\right|}, \quad \cos(\pi - \theta^{(3)}) = \frac{\left(\vec{r}_i^{\,(2)} - \vec{r}^{\,*}\right)\cdot\left(\vec{r}_k^{\,(2)} - \vec{r}^{\,*}\right)}{\left|\vec{r}_i^{\,(2)} - \vec{r}^{\,*}\right|\left|\vec{r}_k^{\,(2)} - \vec{r}^{\,*}\right|} \tag{S2}$$

The fusion rate constant does not depend on the bending angle at the pre-existing degree-2 node.

When fission of a degree-3 junction occurs, an edge is randomly chosen to break off from the others and the junction node is transformed into two separate nodes. The spherocylindrical tip of the broken-off edge is placed at the original junction position, while the new degree-1 node is inset along its edge by a distance $r_s$. The new degree-2 node is shifted away by a distance $r_s$ in the direction opposite to the disconnected edge, slightly compressing the two remaining connected edges (Fig. S1D). This approach ensures that the two new nodes are placed just outside steric contact upon fission. However, it leaves the two remaining connected edges with a small compressive strain that pushes the junction back towards the released node, making it more difficult for the latter to escape prior to re-fusion.

We note that the actual mechanical details and mitochondrial orientations following a fission event are not well-established, and likely involve more complicated interactions with other cellular structures such as tubules of the endoplasmic reticulum and actin filaments [1, 2]. Here, our goal is not to directly represent these details but rather to provide a simplified model for fusion and fission that takes mitochondrial unit orientation into account and does not give rise to extreme mechanical forces following a fusion or fission event.

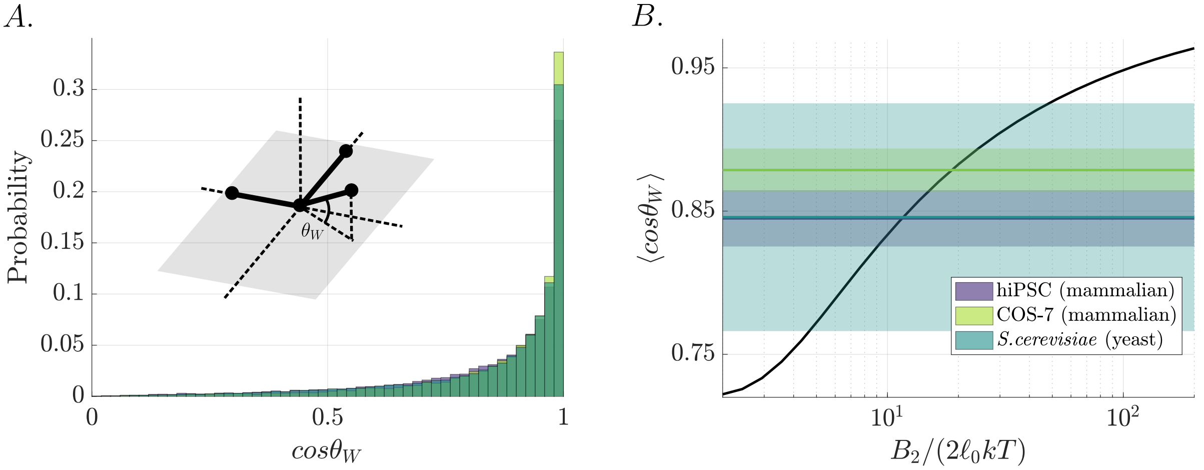


FIG. S2. Selection of the degree-3 bending modulus based on analysis of experimental data. **A.** The distribution of Wilson-type out-of-plane coordinates for all degree-3 junctions is displayed as a histogram for 3 cell types (colors). The distributions largely overlap, even when comparing yeast and mammalian cells. Inset: diagram indicating the out-of-plane coordinate $\theta_W$ for a free degree-3 junction. **B.** Computed average of $\cos\theta_W$ for a free degree-3 junction in our mechanical model (black curve), as a function of junction bending modulus $B_2$. Colored regions indicate mean ± standard deviation across individual cell averages for experimental data from three cell types (as shown in A).

#### S2. SELECTION OF DEFAULT BENDING STIFFNESS AND FUSION SENSITIVITY PARAMETERS $(B_1, B_2, \alpha_1, \alpha_2)$

In our model, $B_1, B_2$ control the bending stiffness at degree-2 and 3 junctions while $\alpha_1, \alpha_2$ control the orientational sensitivity of the fusion machinery at those junctions. To estimate $B_1, B_2$, we consider two experimetally accessible metrics: persistence length for degree-2 junctions and spring stiffness of a Wilson-type out-of-plane coordinate potential [3] for degree-3 junctions. The persistence length of mitochondrial tubules has been previously measured at $\ell_p \approx 2\mu$m by [4] in *Xenopus laevis*. We therefore use $B_1/k_bT^* = \ell_p = 2\mu$m in our simulations unless stated otherwise (with $k_bT^* = 1$ the default temperature). In Fig. 6, where the effective temperature for diffusive motion is altered, the bending modulus is held constant, so that higher motilities correspond to higher average bending angles between mitochondrial segments.

To obtain an estimate for $B_2$, we examined all degree-3 junctions in each of the quantified mammalian network structures from hiPSC and COS-7 cells, as well as the yeast network structures. At each junction, we averaged the three deviation angles of one edge with respect to the plane formed by the other two edges (Fig. S2A), defining the Wilson out-of-plane coordinate, $\theta_W$, which was developed to quantify 3-way junction bending for molecular structures [3]. To relate the bending modulus $B_2$ to this coordinate, we calculate its average for a single thermally fluctuating triskelion according to:

$$\begin{aligned}\langle \cos\theta_W \rangle =& \frac{1}{Z}\int_{-1}^{1} d(\cos\theta^{(1)}) \int_{-1}^{1} d(\cos\theta^{(2)}) \int_{0}^{2\pi} d\phi \, \sqrt{1-(\sin\theta^{(1)}\sin\phi)^2}\; e^{-E_{\text{bend},2}(\theta^{(1)},\theta^{(2)},\phi;B_2)/k_bT^*} \\ Z =& \int_{-1}^{1} d(\cos\theta^{(1)}) \int_{-1}^{1} d(\cos\theta^{(2)}) \int_{0}^{2\pi} d\phi \;\; e^{-E_{\text{bend},2}(\theta^{(1)},\theta^{(2)},\phi;B_2)/k_bT^*}\end{aligned} \tag{S3}$$

where edge 3 of the triskelion is assumed to lie along the z axis and the spherical coordinates for edge 1 are $(\theta^{(1)}, 0)$ and for edge 2 are $(\theta^{(2)}, \phi)$.

The expected value $\langle \cos\theta_W \rangle$ is plotted in Fig. S2B as a function of the bending modulus, with stiffer modulus resulting in less out-of-plane deviation. We compare these results to the measured average for mammalian (hiPSC and COS-7) and yeast cell mitochondrial networks, obtaining a corresponding range of dimensionless bending modulus values $\frac{1}{2\ell_0 k_b T^*} B_2 \approx 10 - 20$. We use a default value of $\frac{1}{2\ell_0 k_b T^*} B_2 = 10$ throughout our simulations.

By default, we set the angular sensitivies $\alpha_1, \alpha_2$ equal to the bending moduli $B_1, B_2$ via the conversions: $\alpha_i = B_i/\ell_0 k_b T^*$, giving the dimensionless $\alpha_1 = 4, \alpha_2 = 20$. For tip-tip interactions, this choice allows the distribution of successful fusion angles (controlled by $\alpha_1$) to approximately match the distribution of fission angles (controlled by $B_1$) in the well-mixed limit. The equilibrium analogy is only approximate, however, as the distribution of node separations at fusion and fission do not match, and mechanical strain is induced following fission. For tip-side fusion, the angular sensitivity only considers the two angles involving the incoming edge, so that the the remaining degree-2 junction is left more bent after a fission event than it would typically be a upon fusion. This deviation from equilibrium is in part responsible for the super-linear dependence of the association constant on the microscopic tip-side fusion rate $k_{u2}$.

#### S3. ESTIMATING ESCAPE PROBABILITIES FOR TWO NODES IN CONTACT

Here we derive estimates for the correction factors $p^*_{\text{esc}}, p_{\text{esc}}$ that give the approximate probability of two nodes to escape from contact range either immediately after a fission event or after approaching each other from a distance.

We consider a single node originally placed at distance $r_0$ from a potential fusion partner. Define $\epsilon(t)$ to be the energetic component determining the fusion rate $k(t)$ between the two nodes, as described by Eq. 4. This component depends on the angular sensitivity $\alpha_i$ and the orientations of the segments involved. We assume that during the time interval before the node either escapes or fuses, the energetic component transitions from its original value $\epsilon_0$ to an equilibrated value $\epsilon_{\text{eq}}$, over a timescale $\tau$. The fusion rate can thus be estimated as

$$k_{ui}^{\text{eff}}(t) = k_{ui} \exp\left[\epsilon_0 e^{-t/\tau} + \epsilon_{\text{eq}}(1 - e^{-t/\tau})\right] \tag{S4}$$

Using the standard separation of variables for a reaction-diffusion system in a spherical domain [5], one can compute the distribution for a particle initially at position $r_0$, diffusing with diffusivity $D_{\text{rel}}$ between a reflecting boundary at $a = 2r_s$ and an absorbing boundary at $b = 2r_c$, subject to time-dependent reaction rate $k(t) = k_{ui}^{\text{eff}}(t)$. Specifically, the survival probability for this particle is given by [5]

$$S(t|r_0, \epsilon_0) = \sum_{m=1}^{\infty} \frac{b}{\mathcal{N}_m \beta_m} R_m(r_0) e^{-D_{\text{rel}}\beta_m^2 t - \int_0^t k(t')dt'}, \tag{S5a}$$

$$\tan \beta_m(b - a) + \beta_m a = 0 \tag{S5b}$$

$$R_m(r) = \sin\left[\beta_m(b - r)\right]/r \tag{S5c}$$

$$\mathcal{N}_m = \frac{1}{2}\left[b - a + a/(1 + \beta_m^2 a^2)\right] \tag{S5d}$$

where $\beta_m, R_m$ are the eigenvalues and eigenfunctions for homogeneous diffusion in a hollow sphere with the desired boundary conditions, and $\mathcal{N}_m$ is a normalization constant. The escape probability is then computed as the probability that the particle does not fuse before hitting the absorbing boundary:

$$p(r_0, \epsilon_0) = 1 - \int_0^{\infty} k(t)\ S(t|r_0, \epsilon_0) dt \tag{S6}$$

We next consider an alternate state where the separation and edge orientation of nearby nodes is proportional to the fusion rate. This is an approximation for state $n$ in the diagram of Fig. 3B, and would give a system that obeys detailed balance if the state $n^*$ were removed. In this case, the initial separation of the two nodes is assumed to be uniform between $a$ and $b$. The escape probability is then computed as:

$$\bar{p}(\epsilon_0) = \frac{3}{b^3 - a^3} \int_a^b r_0^2\ p(r_0, \epsilon_0)\ dr_0 \tag{S7}$$

The appropriate weighted averages over the initial fusion energy component $\epsilon_0$ to calculate $p_{\text{esc}}$ and $p^*_{\text{esc}}$, as well as the relative node diffusivity $D_{\text{rel}}$ and orientation relaxation time $\tau$ are discussed below.

##### Degree-2 junction

For a newly fissed degree-2 junction, we consider the dynamics of the inset nodes, initially separated at $r_0 = 2r_s$. The escape probability is averaged over starting energies $\epsilon_0$, weighed by the Boltzmann factor for the configuration of the degree-2 junction prior to fission ($w_1$). The long-time energetic component is averaged with uniform weighting of angles between the separated segments. Specifically, the weights and averages are defined as:

$$w_1(\theta) = e^{B_1/(\ell_0 k_b T)\cos\theta}, \tag{S8a}$$

$$\epsilon(\theta) = \alpha_1(1 - \cos\theta) \tag{S8b}$$

$$\epsilon_{\text{eq}} = \frac{1}{2}\left[\int_{-1}^{1} \mathrm{d}\!\cos\theta\ \epsilon(\theta)\right] = \alpha_1 \tag{S8c}$$

$$p^*_{\text{esc}} = \left[\int \mathrm{d}\!\cos\theta\ p[2r_s, \epsilon(\theta)] w_1(\theta)\right] / \left[\int \mathrm{d}\!\cos\theta\ w_1(\theta)\right], \tag{S8d}$$

To compute the escape probability for nodes in an equilibrated system, the initial orientations are instead weighed in proportion to the angle-dependent fusion rate, giving:

$$p_{\text{esc}} = \left[\int \mathrm{d}\!\cos\theta\ \bar{p}[\epsilon(\theta)] e^{-\epsilon(\theta)}\right] / \left[\int \mathrm{d}\!\cos\theta\ e^{-\epsilon(\theta)}\right]. \tag{S9}$$

To estimate the relative diffusivity for the two separating nodes ($D_{\text{rel}}$), we consider two 'dumbbells' defined by coordinates $\vec{r_1}, \vec{r_2}$ and $\vec{r_3}, \vec{r_4}$, respectively and subject to the constraints: $f_1(\vec{r_1}, \vec{r_2}) = |\vec{r_1} - \vec{r_2}| - (\ell_0 - r_s) = 0$, $f_2(\vec{r_3}, \vec{r_4}) = |\vec{r_3} - \vec{r_4}| - (\ell_0 - r_s) = 0$. The two dumbbells are assumed to be initially aligned along the z-axis. In one time-step $\Delta t$, each of the dumbbell coordinates moves according to:

$$\Delta\vec{r_i} = \delta\vec{r_i} + \lambda_1 \partial_{\vec{r_i}} f_1 + \lambda_2 \partial_{\vec{r_i}} f_2, \tag{S10}$$

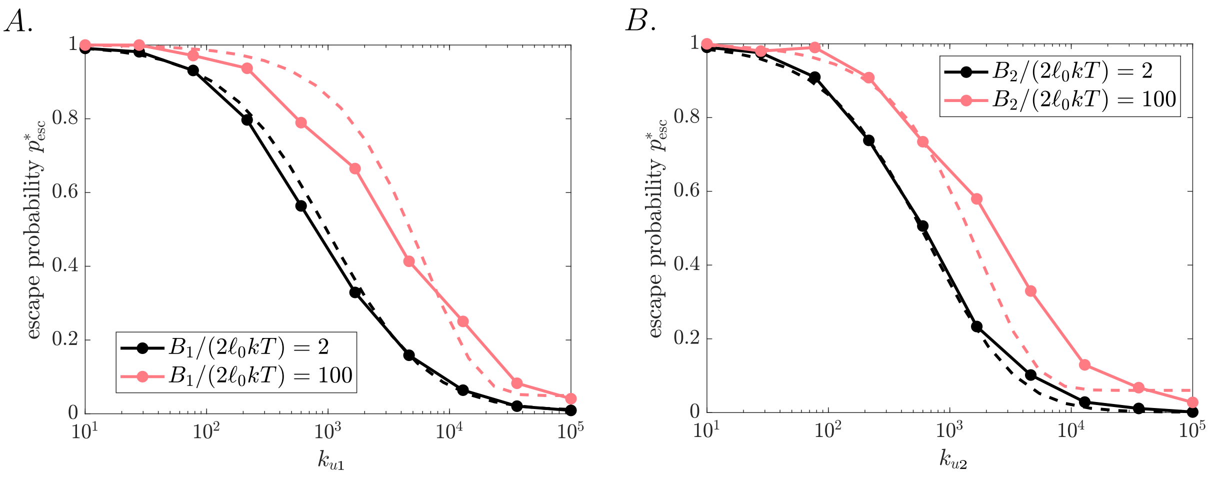


FIG. S3. Computed escape probabilities agree with simulated escape. **A-B.** The non-equilibrium escape probability $p^*_{\rm esc}$ is plotted for the case of a degree-2 (degree-3) junction immediately post-fission as a function of the local fusion parameter $k_{u1}(k_{u2})$. Color indicates the particular value of $B_1(B_2)$ with $\alpha_i = B_i/(\ell_0 k_b T)$. Solid curves are from simulations of isolated pairs of newly-fissed nodes, while dashed curves show the approximate $p^*_{\rm esc}$ predictions.

where $\delta\vec{r}_i$ is the shift due to the Brownian forces, such that $\langle \delta r_{ia} \delta r_{jb} \rangle = 2(k_b T/\mu)\Delta t \; \delta_{ij}\delta_{ab}$. The $\lambda_i$ are Lagrange multipliers which can be found by enforcing the constraints $f_i = 0$ after the step. Solving for these multipliers, and expanding to lowest order in the $\delta\vec{r}_i$ gives:

$$\left\langle |\Delta\vec{r}_i|^2 \right\rangle = 5\frac{k_b T}{\mu}\Delta t. \tag{S11}$$

From there, the effective relative diffusivity of the two separating nodes is found to be $D_{\rm rel} = \frac{5}{3}\frac{k_b T}{\mu}$.

Similarly, the decorrelation time for the angle between the two dumbbells can be found by expanding Eq. S10 for small perturbations. After a single timestep $\Delta t$, the angle $\theta$ between $\vec{r}_2 - \vec{r}_1$ and $\vec{r}_4 - \vec{r}_3$ has the average:

$$\langle \cos\theta \rangle = 1 - \frac{8}{\ell_0^2}\frac{k_b T}{\mu}\Delta t, \tag{S12}$$

yielding the relaxation timescale $\tau = \frac{\mu \ell_0^2}{8 k_b T}$.

The escape probabilities $p^*_{\rm esc}$ are compared directly to simulations wherein isolated degree-2 junctions are allowed to undergo fission, and we track the probability of the newly separated nodes escaping the contact radius prior to re-fusing. The simplified estimates provided here approximately match the simulations (Fig. S3A).

##### Degree-3 junction

For a newly fissed degree-3 junction, we take an analogous approach, with numerical integrals over the orientations of the edges involved in the reaction. The escape probability $p^*_{\rm esc}$ after a fission event is computed from Eq. S6, by integrating over orientations weighted by a Boltzmann factor $(w_3)$ for the configuration of the 3-way junction prior to fission. The long-time equilibrated energy $\epsilon_{\rm eq}$ is obtained by averaging with a weight corresponding to the energy of the remaining fused pair of segments ($w_2$). The equilibrated escape probability is weighted by the fusion rate. Overall, the weights and averages can be summarized as:

$$w_2(\theta^{(1)}) = e^{B_1/(\ell_0 k_b T)\cos\theta^{(1)}}, \tag{S13a}$$

$$w_3(\theta^{(1)},\theta^{(2)},\theta^{(3)}) = e^{-E_{\text{bend}}(B_2,\theta^{(1)},\theta^{(2)},\theta^{(3)})/k_b T}, \tag{S13b}$$

$$\epsilon(\theta^{(2)},\theta^{(3)}) = \alpha_2[2-\cos(\theta^{(2)}-\pi/3)-\cos(\theta^{(3)}-\pi/3)] \tag{S13c}$$

$$\epsilon_{\text{eq}} = \left[\int \mathrm{d}\Omega\ \epsilon(\theta^{(2)},\theta^{(3)})w_2(\theta^{(1)})\right] / \left[\int \mathrm{d}\Omega\ w_2(\theta^{(1)})\right], \tag{S13d}$$

$$p^*_{\text{esc}} = \left[\int \mathrm{d}\Omega\ p(2r_s,\epsilon(\Omega))w_3(\Omega)\right] / \left[\int \mathrm{d}\Omega\ w_3(\Omega)\right], \tag{S13e}$$

$$p_{\text{esc}} = \left[\int \mathrm{d}\Omega\ \bar{p}(\epsilon(\Omega))e^{-\epsilon(\Omega)}\right] / \left[\int \mathrm{d}\Omega\ e^{-\epsilon(\Omega)}\right], \tag{S13f}$$

where orientation $\Omega$ describes the orientations of all three interacting segments, defining the angle $\theta^{(1)}$ between the remaining fused segments and the angles $\theta^{(2)},\theta^{(3)}$ with the newly fused segment. Specifically, the orientations of the two remaining fused segments are taken to be at $(0,0,1)$ and $(\sin\theta^{(1)},0,\cos\theta^{(1)})$. The newly fissed node is placed at position $(-\frac{3}{4}(b^4-a^4)/(b^3-a^3),0,0)$ (corresponding to uniformly averaged separation), and the orientation of the newly fissed segment is defined to be $(\sin\theta^{(2)}\cos\phi,\sin\theta^{(2)}\sin\phi,\cos\theta^{(2)})$. To approximately account for sterically excluded configurations, the azimuthal angle is limited to be above $\phi^* = \tan^{-1}(2r_s/(r_s+r_c))$. The orientational integral is then given by $\int \mathrm{d}\Omega = \int_{-1}^{1}\mathrm{d}(\cos\theta^{(1)})\int_{-1}^{1}\mathrm{d}(\cos\theta^{(2)})\int_{\phi^*}^{(2\pi-\phi^*)}\mathrm{d}\phi$.

Using the same approach as in Eq. S10, wherein each node is perturbed by a small Brownian step, and segment lengths are enforced via Langrange multipliers, the mean squared displacement during a timestep $\Delta t$ of the central node in a pair of connected dumbbells is given by

$$\langle|\Delta\vec{r}_2|^2\rangle = \left\{\frac{7}{3}\frac{k_b T}{\mu}\Delta t,\ \ \frac{31}{15}\frac{k_b T}{\mu}\Delta t\right\} \tag{S14}$$

where the first value corresponds to the case where the two edges are initially aligned, and the second value to the case where they are bent at a 60° angle. The relative diffusivity between this central node and a nearby node of a disconnected dumbbell is then $\left\{1.61\frac{k_b T}{\mu}, 1.52\frac{k_b T}{\mu}\right\}$ for the two cases.

The orientation relaxation of the edges after fission involves both rotational diffusion of the fissed single edge and the mechanical relaxation of the bending angle between the two edges that remain connected. We estimate the latter by noting that the rotational friction coefficient for each terminal node in the dimer, around the central node, is $\mu_r = \mu\ell_0^2$. Given that both of the terminal nodes can move to enable the dimer angle to relax, the relaxation timescale can be estimated as half the friction coefficient of each divided by the effective angular spring constant: $\tau_{\text{mech}} \approx \frac{1}{2}\times\frac{\mu_r}{B_1/(2\ell_0)} = \frac{\mu\ell_0^3}{B_1}$. The timescale for reorientation of the fissed single edge is $\tau_{\text{diff}} \approx \frac{\mu\ell_0^2}{4k_b T}$. For the parameters used in this manuscript ($B_1 \geq 2\times(2\ell_0 k_b T)$), the diffusive relaxation is slower, and we thus use $\tau = \tau_{\text{diff}}$, $D_{\text{rel}} = 1.61\frac{k_b T}{\mu}$ for estimating the escape probabilities.

The escape probabilities $p^*_{\text{esc}}$ are compared directly to simulations of isolated junctions undergoing fission and then either re-fusing or escaping the contact range. The results approximately match the approximations described here (Fig. S3 B).

**Mean separation time**

When considering the relevant recharge rate for permitting escape of newly-fissed nodes prior to re-fusion, we calculate the mean first passage time for a particle with diffusivity $D_{\rm rel}$ in a spherical shell with reflecting boundary at $2r_s$ and absorbing boundary at $2r_c$ to leave that outer absorbing boundary. This mean first passage time can be computed as

$$\tau_{\rm esc} = \left(1 + 2\frac{r_s}{r_c}\right)\frac{(2r_c - 2r_s)^2}{6D_{\rm rel}}, \tag{S15}$$

using the Laplace-transformed diffusion equation in spherical coordinates [6]. Plugging in the dimensionless values $r_s = 0.1$, $r_c = 0.15$, $D_{\rm rel} \approx 1.6$ (for both degree-2 and degree-3 fissions), gives an estimated dimensionless escape time of $\tau_{\rm esc} \approx 0.0024$. Consequently, recharge rates with $k_r \lesssim 400$ should be sufficient to allow the nodes to escape from contact prior to re-fusion.

[1] R. G. Abrisch, S. C. Gumbin, B. T. Wisniewski, L. L. Lackner, and G. K. Voeltz, Journal of Cell Biology **219**, e201911122 (2020).
[2] T. S. Fung, R. Chakrabarti, and H. N. Higgs, Nature Reviews Molecular Cell Biology , 1 (2023).
[3] S.-H. Lee, K. Palmo, and S. Krimm, Journal of computational chemistry **20**, 1067 (1999).
[4] A. B. Fernández Casafuz, M. C. De Rossi, and L. Bruno, Scientific Reports **13**, 4065 (2023).
[5] D. W. Hahn and M. N. Özisik, *Heat conduction* (John Wiley & Sons, 2012).
[6] S. Redner, *A Guide to First-Passage Processes* (Cambridge University Press, 2001).
